# Influenza A virus infection during pregnancy increases transfer of maternal bloodborne molecules to fetal tissues

**DOI:** 10.1101/2025.05.27.656420

**Authors:** Rafael J. Gonzalez-Ricon, Ashley M. Otero, Izan Chalen, Jeffrey N. Savas, Adrienne M. Antonson

**Affiliations:** Neuroscience Program, University of Illinois Urbana-Champaign, Urbana, IL, USA; Department of Animal Sciences, University of Illinois Urbana-Champaign, Urbana, IL, USA; Department of Neurology, Northwestern University Feinberg School of Medicine, Chicago, IL, USA

**Keywords:** influenza virus, blood brain barrier, fibrinogen, maternal immune activation, fetal microglia, neurodevelopmental disorders, placenta

## Abstract

Influenza A virus (IAV) infection during pregnancy is linked to heightened risk for neurodevelopmental disorders (NDDs) in the offspring. The precise pathophysiological mechanism(s) underling this association remains an active topic of research. We propose that maternal immune activation (MIA) triggered by IAV infection can disrupt selective permeability at the maternal-fetal interface, leading to increased transfer of blood-derived molecules into the fetal compartment. Some of these molecules might be responsible for the initiation of inflammatory cascades implicated in NDD etiology. Using a murine model of seasonal IAV infection during pregnancy, we examined placental and fetal brain barrier properties following maternal IAV challenge. Our findings demonstrate an enhanced transplacental transfer of fluorescently labeled tracers from maternal circulation to key neurodevelopmental regions, including the subventricular zone (SVZ) and choroid plexus (ChP) of fetal brains. This effect was most pronounced in fetuses from dams exposed to the highest dose of IAV. Notably, a similar pattern was observed for accumulation of the bloodborne neuroinflammatory molecule fibrinogen in these same brain regions, which was further amplified in response to the highest IAV dose. Moreover, fibrinogen accumulation was positively correlated with Iba1^+^ cell immunofluorescence, suggesting a potential interaction between fibrinogen and Iba1^+^ cells. Collectively, these findings suggest that IAV-induced MIA enhances transplacental transfer of blood-derived molecules into fetal tissues, potentially activating proinflammatory pathways in Iba1^+^ cells.

**Highlights:**

- Maternal influenza infection increases fetal exposure to maternally derived tracers.
- Fetal blood brain barrier dysfunction is evident in the SVZ and ChP.
- Fibrinogen accumulation in the SVZ and ChP correlates with Iba1 intensity.
- Increased vascular permeability may contribute to altered fetal brain development.

## 1. Introduction

Epidemiological research indicates an association between maternal exposure to influenza A virus (IAV) during pregnancy and an elevated risk of neurodevelopmental disorders (NDDs) in offspring, including schizophrenia (SZ) and autism spectrum disorders (ASD) (Mednick 1988; Alan S. Brown and Meyer 2018; A. S Brown 2006). The activation of the maternal immune system—referred to as ‘maternal immune activation’ (MIA)—is hypothesized to be a critical factor in disrupting neurodevelopment during gestational infection with non-vertically transmitted viruses. It is postulated that maternal inflammatory mediators, elicited in response to infection, induce maladaptive processes within the developing fetal brain (Kwon, Choi, and Huh 2022; Littauer and Skountzou 2018; Choi et al. 2016; Wu et al. 2017), rather than the virus itself. However, the precise molecular pathways, and the spectrum of inflammatory molecules involved, remain unclear.

During gestation, the fetus is protected from maternal insults via distinct tissue barriers. The placenta—a blastocyst-derived specialized vascular organ—functions as the main maternal-fetal interface, allowing selective molecular crosstalk between maternal and fetal compartments (Robbins et al. 2010; Zeldovich et al. 2013; Cardenas et al. 2010). While the placenta has historically been overlooked in the context of neurodevelopment (Maltepe and Fisher 2015), several studies have shown that it is responsive to maternal inflammation and integral for conducting immune and endocrine signals to the fetus (Goeden et al. 2016; Zengeler et al. 2023; Liong et al. 2020; Wu et al. 2017). We have previously shown that even moderate gestational respiratory IAV infection in mice results in placental lesions, consistent with inflammation-induced malperfusion (Antonson et al. 2021).

The fetal blood-brain barrier (BBB) constitutes another critical checkpoint in safeguarding the developing brain. In mice, the BBB remains incompletely developed until after birth (Daneman et al. 2010), leaving the fetal brain relatively more vulnerable to external insults during gestation (Petrik et al. 2013). Maternally derived pro-inflammatory mediators have been shown to directly regulate aberrant neurodevelopment in murine models of MIA (Ben-Yehuda et al. 2020; Hsiao and Patterson 2011; Kim et al. 2017; Choi et al. 2016; Smith et al. 2007; Wu et al. 2017). However, less is known about the extent to which these mediators may also compromise the nascent BBB (Banks, Kastin, and Gutierrez 1994; Simões, Generoso, et al. 2018). In particular, the role of fibrinogen—a circulating anticoagulant protein synthesized by the liver and abundantly expressed during pregnancy (Cui, Ma, and Qiao 2020; Vilar et al. 2020)—remains unexplored in MIA models.

Fibrinogen is typically absent in the brain parenchyma under healthy physiological conditions (Saitoh, Tanabe, and Muramatsu 2022). However, numerous studies have demonstrated that in the context of neuroinflammatory conditions like multiple sclerosis (MS), traumatic brain injury (TBI), and Alzheimer’s Disease (AD), fibrinogen can breach the BBB and contribute to neuronal damage. This process is primarily mediated through microglial responses, oxidative stress, and subsequent neuronal cell death (Davalos and Akassoglou 2012; Schachtrup et al. 2007a; Ryu et al. 2015; Merlini et al. 2019). Notably, microglia migrate to the developing fetal brain early during gestation (approximately embryonic day [E]9.5-10.5 in the mouse) (Brioschi et al. 2023) and are key cellular mediators of healthy neurodevelopmental processes (Antony et al. 2011; Squarzoni et al. 2014; Tay et al. 2017). While microglia are known to shift their functions during MIA (Otero and Antonson 2022), the extent to which fibrinogen leakage into the developing fetal brain may contribute to these responses has never been explored.

In this study, we hypothesized that IAV-induced MIA disrupts the integrity of placental and fetal brain vascular barriers, enabling the transfer of proinflammatory molecules such as fibrinogen into the developing fetal brain. Using our model of moderate and severe respiratory IAV infection in pregnant mice (Otero et al. 2024), we observed increased transfer of large molecular weight tracers across the maternal-fetal interface, accumulating in the fetal subventricular zone (SVZ) and choroid plexus (ChP)—regions critical for neurodevelopment (H. Xu et al. 2021; Tanabe et al. 2023; Luo and Wang 2024; Eşiyok and Heide 2023). In parallel experiments, we found that fibrinogen also accumulated in these areas and was associated with elevated immunofluorescence of microglia marker, Iba1. These findings suggest that MIA may modulate fetal microglial responses via enhanced transfer of immune-related molecules into the fetal brain, potentially altering normal brain development during embryogenesis.

## 2. Materials and methods

### 2.1. Animals

Nulliparous 9-to-10-week-old female C57BL/6JnTac mice from Taconic Biosciences (Germantown, NY) were acclimated for a minimum of one week at the University of Illinois Urbana-Champaign mouse vivarium before breeding. Females were trio-bred for three to four days. The presence of a vaginal plug was considered gestational day (GD) 0.5. Female body weights were recorded daily until sacrifice. All mice were maintained under a twelve-hour light/dark cycle (7:00 am lights on/ 7:00 pm lights off) and fed ad libitum. A total of 27 pregnant mice across three treatment groups were used to examine maternal and fetal tissues at embryonic (E) day 16.5. A separate cohort of 15 pregnant dams was used to examine tracer distribution in the placenta and fetus. All experimental procedures were approved by the Institutional Animal Care and Use Committee (IACUC) at the University of Illinois Urbana-Champaign.

### 2.2. Influenza A virus inoculation

Mouse-adapted IAV strain X31 (H3N2) was propagated and titered as previously described (Kenney et al. 2019). At E9.5, which corresponds approximately to the end of the first trimester in humans, pregnant mice were randomly assigned to treatment groups and inoculated intranasally with 10^3^ (X31_mod_) or 10^4^ (X31_hi_) tissue culture infectious dose 50 (TCID_50_) IAV, or mock-inoculated with sterile phosphate-buffered saline (Con), under light isoflurane anesthesia as previously described in (Antonson et al. 2021). Maternal and fetal tissues were collected at 7 days post inoculation (dpi).

### 2.3. Tissue collection

Pregnant mice were euthanized under CO_2_ inhalation at 7 dpi, corresponding to the anticipated peak of the inflammatory response against IAV as established by previous studies (Jian Wang et al. 2014; Kenney et al. 2019). Tissues were then collected under sterile conditions. After the uterus was excised, it was promptly transferred to a sterile petri dish containing phosphate-buffered saline (1X PBS; pH= 7.4), and conceptuses were assessed individually. Placental weight, fetal weight, fetal crown-rump length, fetal viability, and reabsorptions were recorded. Individual fetal heads and placentas were either snap-frozen on dry ice or placed in 10% neutral buffered formalin (NBF). Maternal spleen weights were recorded as a marker of systemic inflammation in response to viral infection (Antonson et al. 2021). Maternal lungs were snap-frozen for viral PCR.

### 2.4. Tracer injection and tissue processing

A subset of 15 pregnant dams were retro-orbitally injected with 100 µL of a cocktail of tracers at 7 dpi under anesthesia. One dam per group was injected with 100 µL of sterile PBS to serve as a histological control. For tracer administration, dams from all treatment groups were anesthetized with 4% isoflurane. A drop of ophthalmic anesthetic (0.5% proparacaine hydrochloride ophthalmic solution) was added to both eyes. Next, the animal was injected into the retro-bulbar sinus with 100 µL of tracers or sterile PBS. Mice were allowed to recover from anesthesia and were monitored until tissue collection. The fluorescent tracer mixture consisted of 0.5 mg/mouse of Dextran-Cy5 Fixable (HAWORKS, 250 KDa MW, Catalog no: DE-CyDye-250k), 0.5 mg/mice of Dextran-Texas RedTM (Dextran, Texas RedTM, 70,000 MW, Lysine Fixable, Catalog no: D18464), and 0.5 mg/mice of Dextran-Fluorescein (Dextran, Fluorescein, 500,000 MW, Anionic, Lysine Fixable, Catalog no: D7136). 60 minutes after i.v. injection, whole fetuses and placentas were excised and preserved in 10% NBF, then cryopreserved in 30% sucrose and stored in OCT at −80 °C. Each fetus and placenta were sagitally cryosectioned at 25 µm, mounted on Fisherbrand™ Superfrost™ Plus Microscope Slides (Catalog no. 22-037-246), stained with DAPI for 1min, and stored at 4_°_C until imaging.

### 2.5. Immunohistochemistry

Placentas and fetal heads were fixed in 10% NBF for 24h at 4_°_C and then washed twice in 1x PBS (pH=7.4) for 10 minutes each. Tissues were subsequently immersed in 30% sucrose solution with sodium azide at 4_°_C for 48 h or until the samples sank. Placentas and fetal brains (dissected from heads after fixation) were embedded in OCT tissue-tek (Thermo Fisher Scientific, Catalog no. NC9159334) for cryoprotection and stored long-term at -80_°_C. 10 µm sagittal sections of placentas and fetal brains were cryosectioned and transferred to Fisherbrand™ Superfrost™ Plus Microscope Slides (Catalog no. 22-037-246) for on-slide staining. Sections were washed three times in PBS with 0.05% Tween-20 (Thermo Fisher Scientific, Catalog no. PRH5152; PBST) for 5 min each time and subsequently incubated with blocking buffer (5% Donkey serum [Millipore sigma, Catalog no. 711-585-152], 1% bovine serum albumin [Thermo Fisher Scientific, Catalog no. 126609100GM], 0.3% Triton-X 100 [Thermo Fisher Scientific, Catalog no. ICN19485450] in PBST) for 1 h at room temperature. For fibrinogen staining, donkey serum was omitted from the blocking buffer to reduce background noise. Fetal brain and placental sections were incubated overnight at 4_°_C with primary antibodies. Fetal brain antibodies: rabbit anti-CLDN5 (1:1000; ThermoFisher, Catalog no. 34-1600), goat anti-CD31 (1:250; R&D Systems, Catalog no. AF3628SP). Placental antibodies: rabbit anti-CLDN1 (1:50; ThermoFisher, Catalog no. 51-9000), rabbit anti-CLDN5 (1:300; ThermoFisher, Catalog no. 34-1600), rabbit anti-OCLN (1:100; ThermoFisher, Catalog no. 71-1500), and goat anti-CD31 (1:250; R&D Systems, Catalog no. AF3628SP). For fibrinogen & Iba1 staining in fetal brain sections, sheep anti-fibrinogen (US biological, provided by Professor Katerina Akassoglou Lab, Gladstone Institutes, University of California San Fransisco), sheep anti-fibrinogen (US biological F4203-02F, coagulation factor I), and rabbit anti-Iba1 (1:1000; Wako Chemicals U.S.A, Richmond, VA, Catalog no. 019-19741), were used. Sections were washed three times in PBS-T and incubated with secondary antibodies—Alexa Fluor 594 AffiniPure Donkey Anti-Rabbit IgG H&L (1:1000; Jackson Immuno Research, West Grove, PA, Catalog no. 711-585-152), Alexa Fluor 488 AffiniPure Donkey Anti-goat IgG H&L (1:1000; Jackson ImmunoResearch, Catalog no. 705-545-003), or Alexa Fluor 488 Donkey Anti-Sheep IgG H&L (1:1000; Jackson ImmunoResearch, Catalog no. 713-155-003)—for 2 h followed by staining in DAPI (Thermo Fisher Scientific, Catalog no. EN62248) for 1 min. Sections were mounted with Fluoromount-G Mounting Medium (Thermo Fisher Scientific, Catalog no. 5018788) and stored long-term at 4_°_C.

### 2.6. Imaging & Image Analysis

All images were acquired using a ZEISS AxioScan.Z1 slide scanner at The Carl R. Woese Institute for Genomic Biology (IGB). The configurations were as follows: Colibri 7 LED light source: 385 nm, 430 nm, 511 nm, 555 nm, 590 nm, 630 nm; Objectives: 5x/0.25 10x/0.45, 20x/0.8, 40x/0.5 Pol and 50x/0.8 Pol; Filter: GFP, DsRed, Cy5, DAPI/GFP/CY3/Cy5, and CFP/FP/mCherry; Camera: Hamamatsu Orca Flash, AxioCam IC (CCD AxioCam IC Color camera) and Hitachi HV-F202SCL. Before processing, the images were subjected to shading correction to ensure uniform lighting, followed by blinded analysis using ZEISS ZEN 3.0 Blue software (Oberkochen, DE). Tracer fluorescence intensity was analyzed using ZEISS ZEN 3.0 Blue software. Regions of interest (ROIs)—including the whole fetus, whole placenta, fetal brain (whole SVZ, and ChP) fetal liver, decidua, junctional zone, and labyrinth—were outlined, and fluorescence intensity was quantified for each channel. Negative control samples, obtained from dams injected with sterile saline, were imaged under the same settings and used to establish autofluorescence baselines (**Supp. Fig. S1**). As shown in **Supp. Fig. S1M-O**, the mean fluorescence intensity per area (MFI/mm²) in the fetal liver was the only ROI to exhibit variability across treatment groups. To account for this and to ensure consistency, fetal liver base MFI/mm² (e.g., autofluorescence) was subtracted from all samples across all channels, as depicted in **Supp. Fig. S1Zii**. The remaining ROIs were quantified and expressed as absolute values without autofluorescence subtraction. For quantification of tracer MFI in the fetal SVZ and ChP, fewer of the sections contained a complete cross-section of the ChP; thus, a max total of five fetuses per group were analyzed for the ChP versus ten for the SVZ.

Mean fluorescence of proteins of interest (antibody-based immunofluorescence) was quantified using Image J. Quantification of immunofluorescence within the placenta was performed for each distinct tissue layer—decidua, junctional zone, and labyrinth. Fetal brain Iba1^+^ cells were counted manually and normalized by area (cells/mm^2^). Fibrinogen and Iba1 MFI/mm² values in the ChP were divided by 1,000 to facilitate data visualization and improve graph interpretability, given the relatively high fluorescence intensity measured in this region. Placental and fetal brain ROIs were identified based on reference atlases (Schambra 2008; GENSAT, n.d.; Chen et al. 2017; Allen Institute for Brain Science 2004). The Allen brain atlas (Allen Institute for Brain Science 2004) was utilized to identify the SVZ in fetal brain tissue (sagittal brain images 9-11 at E15.5).

### 2.7. Quantitative real-time PCR

RNA from fetal brains and placentas was isolated using TRIzol via the TRI reagent protocol from Invitrogen (Carlsbad, CA). cDNA was produced using the High-Capacity cDNA reverse transcription kit (Applied Biosystems) following the manufacturer’s instructions, and qPCR was performed on QuantStudio 5 using SYBR Green Master Mix (Applied Biosystems). The 2^−ΔΔCt^ method was used to analyze samples relative to *Rplp0* or *Hprt1* housekeeping genes. **Table S1** contains the list of primers used for gene expression, acquired from Integrated DNA Technologies (IDT) Custom DNA Oligos.

### 2.8. Statistics

Unless otherwise specified, data were analyzed using GraphPad Prism 9 software (San Diego, CA) with significance set at α = 0.05. A randomly selected fetus and placenta from each litter served as the experimental unit. Con, X31_mod_, and X31_hi_ groups were compared using one-way ANOVA assuming Gaussian distribution and homogeneity of variance of residuals, and Tukey correction for multiple comparisons was used. For data with unequal variance, Brown-Forsythe ANOVA was performed with Dunnet T3 correction for multiple comparisons. For non-parametric data, Kruskal-Wallis ANOVA was performed with Dunn’s correction for multiple comparisons. Kruskal-Wallis was used in the case that residuals did not meet normality or homogeneity of variance. Two-way ANOVA (repeated measures) with Tukey post hoc multiple comparisons was used for dam body weight trajectories across gestation.

For litter measurements, all fetuses and placentas in each litter were included. For tracer experiments, an average of five randomly selected fetuses and placentas per litter were included. This allowed us to include the majority of fetuses from smaller litters and get an accurate sampling of fetuses from larger litters. Residual normality was assessed, and due to the non-normal distribution of the data, a natural logarithmic transformation was applied (Hamasha et al. 2022). A constant value of 2 was added to each MFI/mm² measurement before natural logarithmic transformation to account for negative data values (Wicklin Rick 2011). Then, to account for within-litter variability, data were analyzed using a mixed-effects ‘nlme’ model in R with litter as a random effect (Golub and Sobin 2020).

Outliers were identified and removed using the 2 standard deviations rule for gene expression data, or Robust regression and Outlier removal (ROUT) method with a Q=1% for all other analyses. Each statistical test is indicated in the figure legends. All data are expressed as means ± SEM.

## 3. Results

### 3.1. IAV triggers localized and systemic immune responses during gestation

To assess maternal immune responses to moderate and high dose IAV (i.e., X31_mod_ and X31_hi_), the body mass of pregnant dams was recorded daily. In line with our previous findings (Otero et al. 2024; Antonson et al. 2021), body weight gain stagnated in infected groups starting at 4 dpi and lasted to 7 dpi (**Fig. 1A**). Viral RNA was present in maternal lungs from both IAV-treated groups, confirming infection (**Fig. 1B**). Respiratory IAV infection led to an increase of spleen weight relative to body mass, indicative of systemic anti-viral immune responses (Jiang et al. 2021; Lei et al. 2024) (**Fig. 1C**). IAV infection did not impact litter size, reabsorptions, or fetal viability (**Table S2**), which is consistent with our previous studies (Otero et al. 2024). However, when compared to the control group, both X31_mod_ and X31_hi_ groups had reduced placental weight (**Fig. 1D**). Furthermore, while no changes in fetal weight were observed (**Fig. 1E**), a reduction in fetal length (crown to rump measurement) reached significance for the X31_hi_ group (**Fig. 1F**). These findings indicate that maternal respiratory IAV elicits moderate intrauterine growth restriction without fetal loss.

**Figure 1.**
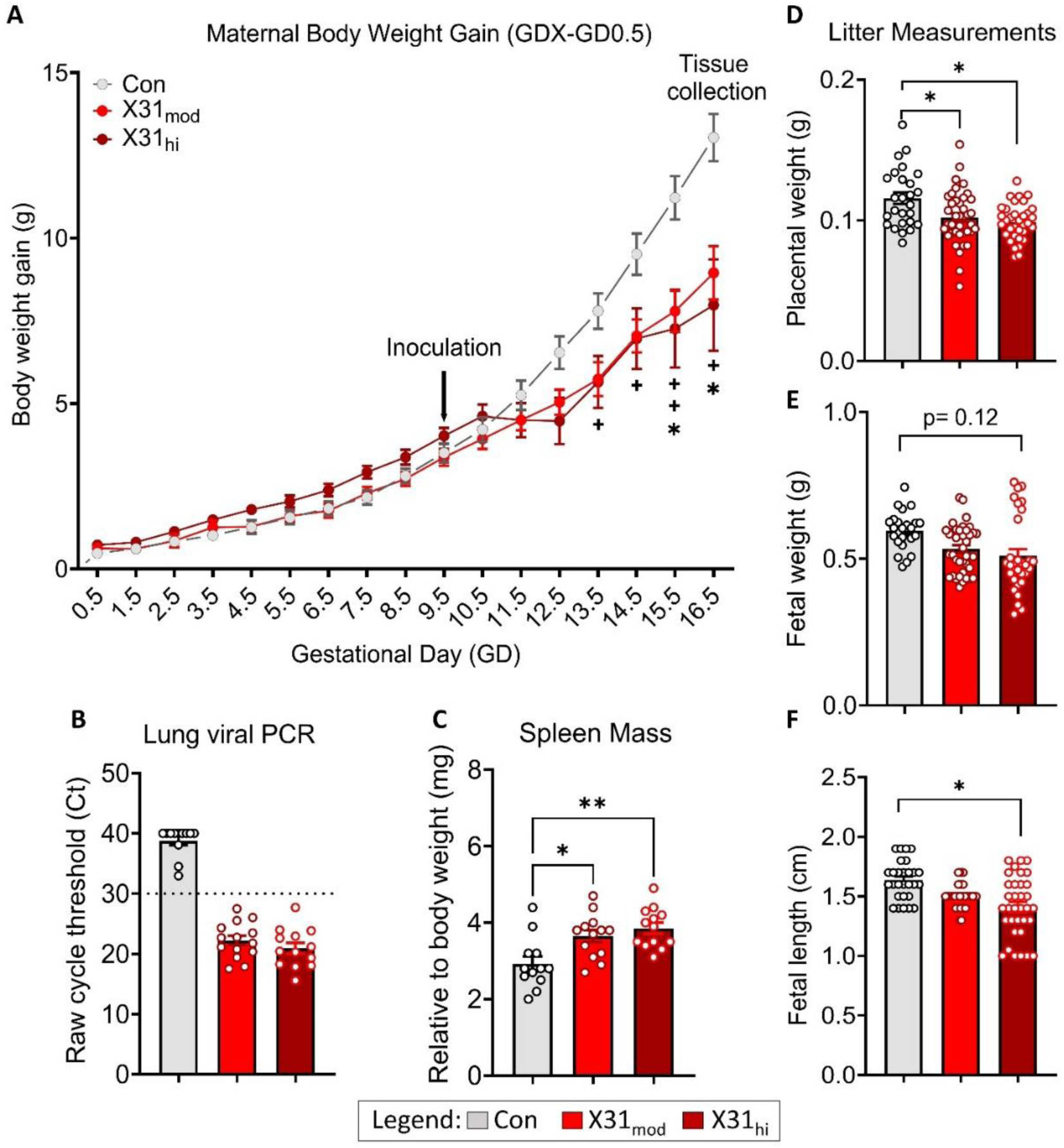
Gestational IAV infection leads to systemic maternal inflammation and reduces placental weight and fetal length. (**A**) Dam body weight gain trajectories (GDX–GD0.5) following IAV inoculation demonstrate a suppression in weight gain in X31_hi_ dams starting at E13.5 and X31_mod_ dams starting at E15.5. Statistical analysis was performed using repeated measures two-way ANOVA with Tukey’s post hoc test for multiple comparisons, revealing a significant main effect of time (p < 0.001).; + = Con vs X31_mod_, * = Con vs X31_hi_; one symbol = p < 0.05, two symbols = p < 0.01. (**B**) Presence of IAV in maternal lungs was confirmed by PCR using a cycle threshold of ≤ 30 cycles (dotted line) as evidence of infection. (**C**) An increase in spleen mass relative to body weight was observed in IAV-infected dams. Placental weight (**D**) and fetal length (**F**) were reduced in X31 litters. (**E**) Differences in fetal weight across groups did not reach significance. In (**C**), groups were compared using a one-way ANOVA with Tukey post hoc multiple comparisons. In (**A-C**), circles represent individual dams; n per group: Con=12- 14, X31_mod_=14-15, X31_hi_=13. In panels (**D-F**), statistical comparisons were performed using a mixed-effects model fitted with the ‘nlme’ package in R, where ‘litter’ was treated as a random effect. Circles represent individual fetuses; n per group: Con = 28, X31_mod_ = 42; X31_hi_ = 34. In (**B-F**), * = p < 0.05, ** = p < 0.01. Non-significant p-values are shown for the comparison of X31_hi_ versus Con. Data are means ± SEM.

### 3.2. Gestational IAV infection increases transfer of high molecular weight tracers across the placenta

During healthy pregnancies, the conceptus is typically shielded from maternal harm by specialized tissue barriers. The placenta serves as the primary physical and immunological wall, blocking the entry of pathogens while selectively permitting the transport of essential nutrients and gases into the fetal blood (Megli and Coyne 2022; Winterhager and Gellhaus 2017). To test whether IAV infection increases transfer of maternally derived molecules into the fetal compartment, a cocktail of fluorescent tracers of varying molecular weights (e.g., TxRed-70 KDa, Cy5-250 KDa, and Fluorescein-500 KDa) was administered intravenously (i.v.) to pregnant dams at E16.5. This set of tracers was selected to approximate the transport dynamics of albumin (66.2 KDa), the most abundant protein in circulation (Murano, Wiman, and Blombäck 1972), as well as fibrinogen (340 KDa), which is especially abundant during pregnancy (Merlot, Kalinowski, and Richardson 2014; Hale et al. 2012). Additionally, larger molecules that typically do not traverse the placental barrier were represented by Fluorescein-500 KDa tracer to evaluate size-restricted transfer across the placenta (Kuna et al. 2018; Dilworth and Sibley 2013; Menjoge et al. 2011; Gupta and Gupta 2017). The procedure for selecting ROIs (i.e., placenta, whole fetus, and fetal brain) is outlined in **Fig. 2A**, and representative images in **Fig. 2E** illustrate the distribution of all tracers across the experimental groups (adjusted for autofluorescence using tracer-free control samples; **Supp. Fig. S1**). A dose-dependent increase in the accumulation of all tracers within whole fetuses was seen in IAV-treated groups (**Fig. 2B**-**D**).

**Figure 2.**
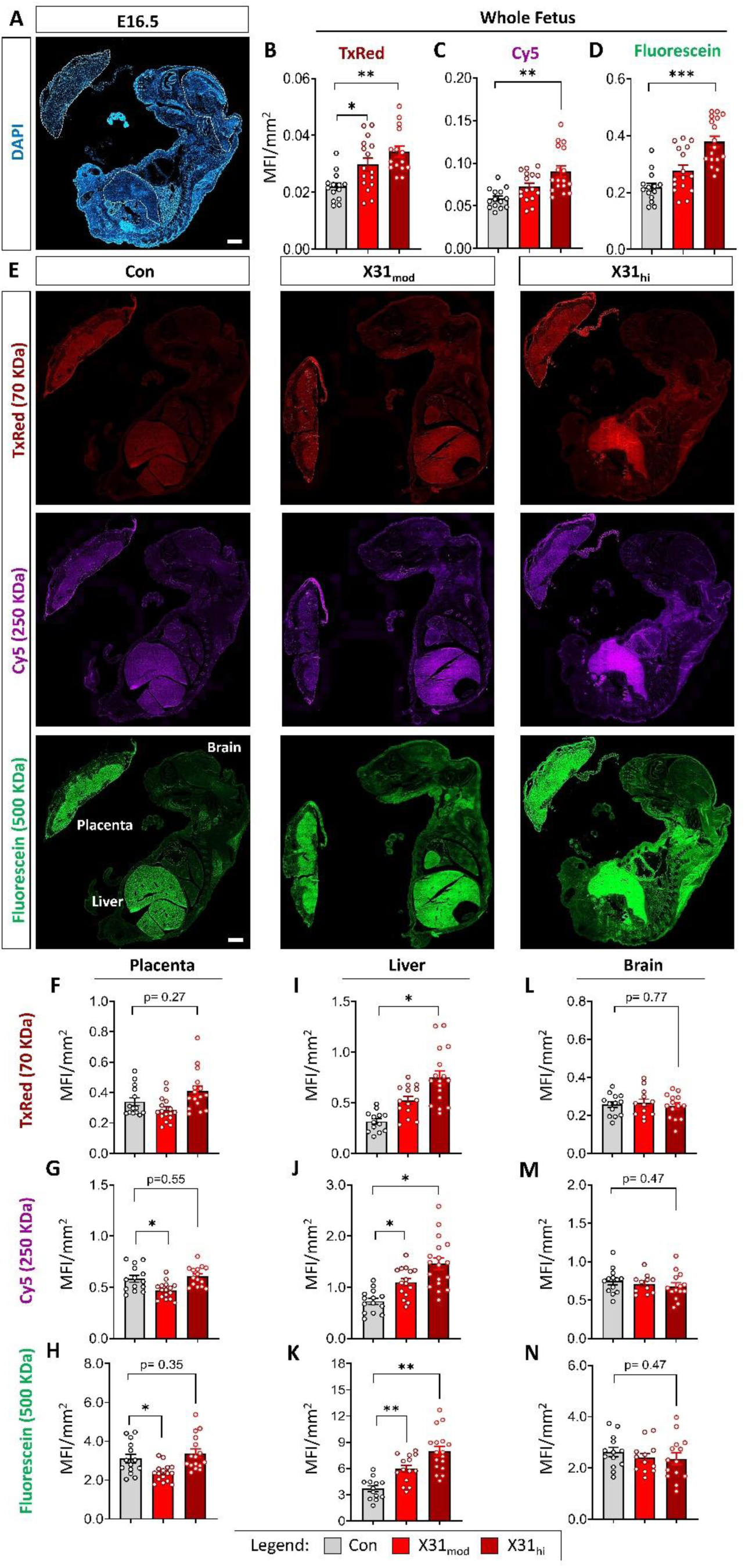
Gestational IAV dose-dependently increases the transfer of maternally derived large molecules into fetal organs at E16.5. (**A**) Representative DAPI image demonstrating the standardized procedure for selecting regions of interest: placenta, fetal brain, and fetal liver. (**B-D**) Prenatal IAV infection led to increased maternal-to-fetal transfer of all tracers in the whole fetus. (**E**) Representative fluorescent cross-sections of fetuses and placentas demonstrating accumulation of different molecular weight tracers across treatment groups, with background autofluorescence subtracted. In the placenta, no differences were observed in TxRed (**F**), while Cy5 (**G**), and Fluorescein (**H**) fluorescence decreased only in the X31_mod_ group. (**I**) A pronounced accumulation of TxRed was observed in the liver of X31_hi_ fetuses, while Cy5 (**J**) and Fluorescein (**K**) accumulated in the liver of both X31_mod_ and X31_hi_ fetuses. (**L-N**) No changes in tracer distribution were seen in fetal brains. P-values are shown for the comparison of X31_hi_ versus Con. Statistical comparisons were performed using a mixed-effects model fitted with the ‘nlme’ package in R, with litter as a random effect; * p < 0.05, ** p < 0.01, *** p < 0.001. Data are expressed as means ± SEM; circles represent individual fetuses; n per group: Con = 13-14, X31_mod_ = 13-16; X31_hi_ = 13-17. Scale bar = 1 mm.

When the whole placenta was quantified, tracer accumulation patterns were more complex. While TxRed MFI/mm^2^ was unchanged (**Fig. 2F**), a drop in Cy5 (**Fig. 2G**) and Fluorescein (**Fig. 2H**) MFI was observed in only X31_mod_ placentas. To investigate how differential tracer accumulation within distinct placental layers might contribute to these findings, we compared fluorescence intensity across the decidua, junctional zone, and labyrinth (**Supp. Fig. S2**). At the decidua, no changes were detected in the MFI/mm^2^ for any of the three tracers (**Supp. Fig. S2A**–**C**). However, the X31_hi_ placentas exhibited increased deposition of TxRed (**Supp. Fig. S2D**) and fluorescein (**Supp. Fig. S2F**) within the junctional zone, with Cy5 MFI/mm² not reaching significance (p = 0.06, **Supp. Fig. S2E**). In the labyrinth, the most vascularized region of the placenta, Cy5, and Fluorescein MFI were reduced exclusively in the X31_mod_ group compared to controls (**Supp. Fig. S2G-I**), contributing to the overall differences observed when the whole placenta is quantified on aggregate. Overall, these findings suggest that IAV-induced MIA influences the transplacental transfer of fluorescent tracers, with varying patterns of distribution across placental layers, particularly in the junctional zone and labyrinth.

Next, we analyzed tracer accumulation in the fetal liver and fetal brain. The fetal liver plays a crucial role in prenatal development by serving as the primary site of embryonic hematopoiesis (Nierhoff et al. 2005; Lewis, Yoshimoto, and Takebe 2021; Yokomizo et al. 2022; Si-Tayeb, Lemaigre, and Duncan 2010). Beyond its hematopoietic role, the fetal liver is essential for the synthesis of a wide range of plasmatic proteins, including fibrinogen, which reach adult levels by E16 (Pickart and Michael Thaler 1979). We observed an increased accumulation of TxRed-70 KDa exclusively within the X31_hi_ group (**Fig. 2I**), followed by the accumulation of Cy5-250 KDa (**Fig. 2J**) and Fluorescein-500 KDa (**Fig. 2K**) on both IAV-treated groups when compared to the control. However, unlike the fetal liver, tracer distribution in fetal brains remained unchanged (**Fig. 2L-N**).

Our findings demonstrate that IAV-induced MIA significantly enhances the transfer of molecules up to 500 KDa from the maternal circulation to the fetus, with predominant accumulation in the fetal liver. This highlights the liver’s key role in metabolizing bloodborne molecules from both maternal and fetal sources (L. D. Brown et al. 2024; Bowman, Arany, and Wolfgang 2021; Bowman et al. 2019). Tracer accumulation was altered within the placenta, with reduced levels in the labyrinth and an increase in the junctional zone, which aligns with previous observations of placental malperfusion following IAV infection (Liong et al. 2020; Antonson et al. 2021). These results emphasize the distinct functional properties of each placental layer in regulating molecular permeability, highlighting their dose-dependent differential responses to maternal anti-viral inflammation.

### 3.3. High molecular weight tracers accumulate in the fetal SVZ and ChP following exposure of dams to high-dose IAV

While the accumulation of fluorescent tracers in the fetal brain did not differ when the brain was quantified as a whole, we predicted that tracers may preferentially accumulate in regions with more permeable barriers. Thus, we conducted a more detailed analysis to determine the extent of tracer penetration at the SVZ and ChP, two regions critical for neurological development linked through the exchange of molecules between the blood and cerebral spinal fluid (CSF) (Naffaa et al. 2023; Arnò et al. 2014; Hattori 2022; Saunders et al. 2023). As shown in **Fig. 3A**, representative images illustrate the distribution of all tracers across groups, with DAPI used as a reference to delineate the regions of interest at the lateral ventricle. While no significant differences were observed in the MFI of the relatively low (TxRed) and medium (Cy5) molecular weight tracers in the SVZ (**Fig. 3B**-**C**) and ChP (**Fig. 3F**-**G**), we detected increased deposition of the highest molecular weight tracer in both regions (**Fig. 3D** & **H**). This increase occurred exclusively in response to the high infectious dose of influenza A virus. This finding suggests that severe maternal immune activation may compromise barrier integrity in these regions, allowing the passage of molecules that are typically restricted.

**Figure 3.**
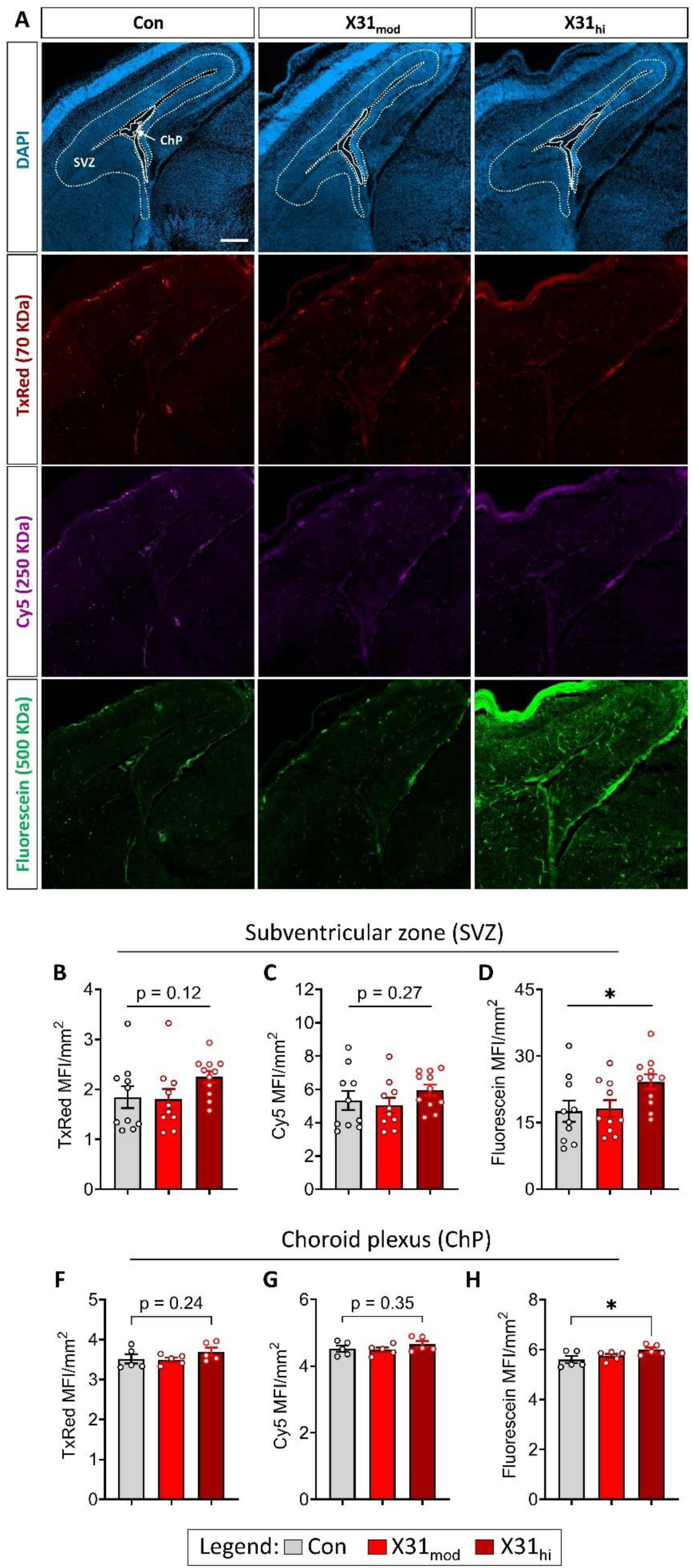
Only X31_hi_ infection enhanced the transfer of fluorescein (500 KDa) from maternal circulation into the fetal SVZ and ChP. (**A, top row**) ROIs were standardized based on DAPI staining, and representative MFI/mm² quantifications of tracers within the SVZ and ChP (**A, bottom rows**) are shown across groups. While no changes were observed for TxRed (**B, F**) or Cy5 (**C, G**), MFI/mm² of fluorescein (**D, H**) increased following only X31_hi_ exposure in both the SVZ and ChP. Data underwent logarithmic transformation (log(x + 2)) before analysis. P-values are shown for the comparison of X31_hi_ versus Con. Statistical comparisons were performed using a mixed-effects model fitted with the ‘nlme’ package in R, with litter as a random effect; * p < 0.05. Data are expressed as means ± SEM; circles represent individual fetuses; n per group: Con (SVZ: n=10, ChP: n=5), X31_mod_ (SVZ: n=10, ChP: n=5), X31_hi_ (SVZ: n=10, ChP: n=5). Scale bar = 250 µm.

### 3.4. IAV exposure does not affect tight junction protein levels in the placenta

MIA induced by IAV can lead to alterations in placental architecture and functionality that may exert aberrant effects on fetal development due to placental vascular dysfunction (Creisher et al. 2024; Antonson et al. 2021; Liong et al. 2020). To assess whether increased tracer dissemination within the fetal compartment correlates with a breakdown in placental vascular endothelial barriers, we analyzed gene expression and immunofluorescence of key tight junction proteins (TJP), including claudins 1-5 (CLDN), zonula occludens-1 (ZO-1), and occludin (OCLN), which are essential for placental barrier integrity (Liévano et al. 2006) (**Fig. 4 & Supp. Fig. S3**). Additionally, we examined immunohistochemical expression of endothelial cell pan marker CD31, which regulates cellular trafficking and vascular permeability (Lertkiatmongkol et al. 2016). Representative images of CLDN5, CD31, CLDN1, and OCLN immunostaining in placental layers are shown in **Fig. 4A** & **Supp. Fig. S3A**, respectively.

**Figure 4.**
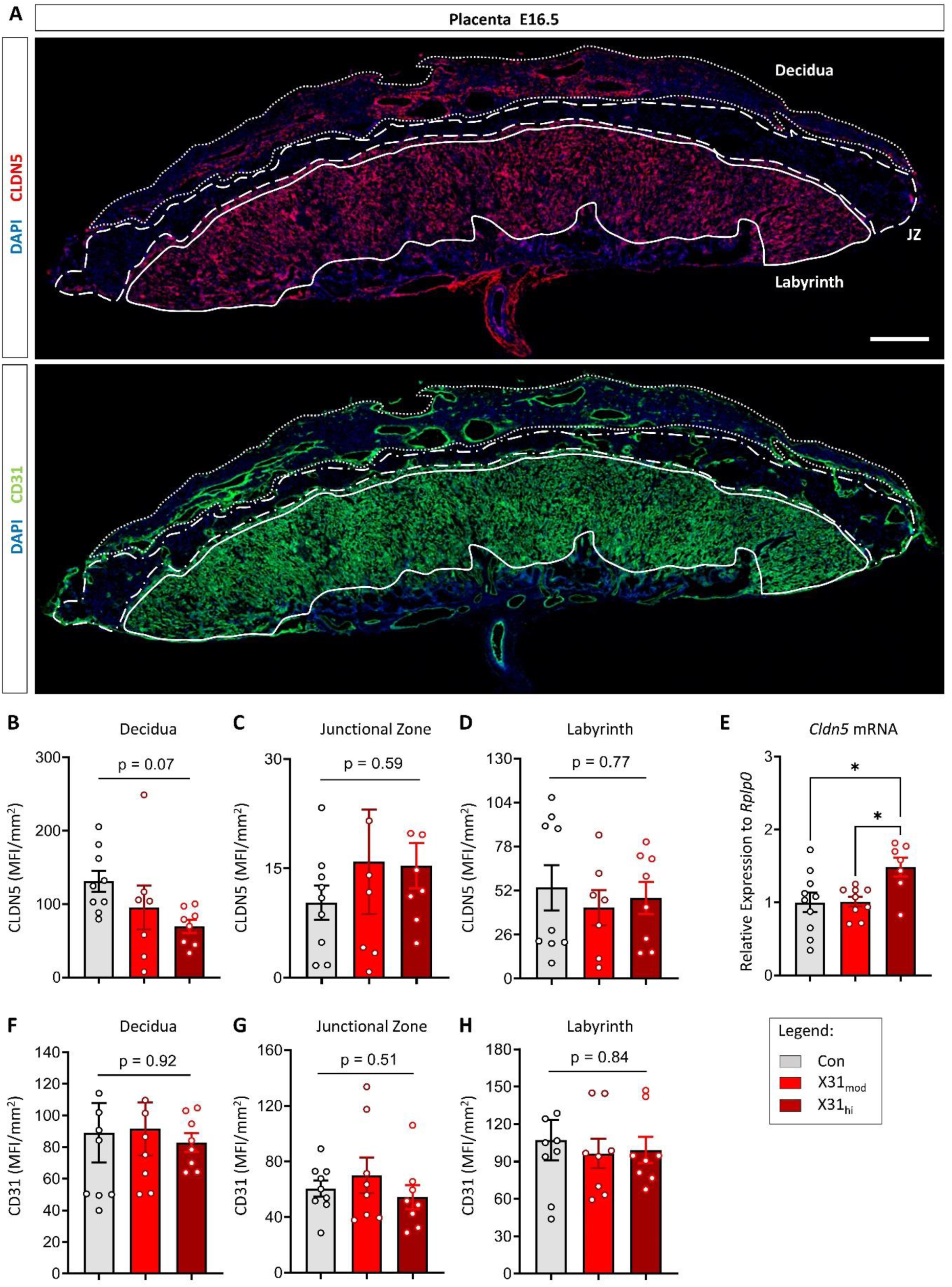
Gestational IAV infection impacts *Cldn5* gene expression in whole placental tissue in a dose-dependent manner. (**A**) Representative cross-sections of CLDN5 and CD31 staining in placental samples at E16.5; dotted, hatched, and solid lines delineate the decidua, junctional zone, and labyrinth, respectively. No differences were seen in (**B-D**) CLDN5 or (**F-H**) CD31 MFI/mm^2^ across placental layers. (**E**) An increase in *Cldn5* gene expression in the whole placenta was observed in only the X31_hi_ group. One representative fetus per litter was used to generate a litter mean for each group. Data are expressed as means ± SEM; circles represent individual placentas; n per group: Con = 8-10, X31_mod_ = 6-9, X31_hi_ = 6-8. Statistical analysis was conducted using a one-way ANOVA with Tukey’s post hoc test; * = p < 0.05. Non-significant p-values indicate main treatment effects (not post hoc). Scale bar = 200 μm.

CLDN5 was the only TJP that exhibited a downward trend, and only in the decidua, with the lowest values in the X31_hi_ group; however, this difference did not reach statistical significance (**Fig. 4B**; p = 0.07). Beyond this trend, no significant differences were observed in the MFI/mm² of CLDN5 across other placental layers (**Fig. 4C**-**D**). Similarly, MFI/mm² of CD31 did not differ across treatment groups (**Fig. 4F**-**H**), indicating overall abundance of vascular endothelial cells was unaffected. Both MFI/mm² of CLDN1 (**Supp. Fig. S3B**-**D**) and of OCLN (**Supp. Fig. S3E-F**) also remained constant. In line with these findings, expression profiles of *Cldn1*, *Cldn2*, *Cldn4*, *Ocln*, and *Zo-1* transcripts were unchanged between groups (**Supp. Fig. S3H-L**). Interestingly, and in contrast to the other TJP transcripts, the expression of *Cldn5* was elevated only in the X31_hi_ group relative to control (**Fig. 4E**).

We then evaluated the expression profile of nitric oxide synthase 2 (*Nos-2*) and cyclooxygenase 2 (*Cox-2*), two key regulators of reactive oxygen species (ROS) synthesis and trophoblast apoptosis (Yuan et al. 2006). However, neither *Nos-2* (**Supp. Fig. S3M**) nor *Cox-2* (**Supp. Fig. S3N**) mRNA levels differed between treatment groups. Taken together, these data indicate that overt changes in endothelial TJPs and oxidative stress are absent in the placenta following maternal IAV exposure, despite clear evidence of increased transplacental transfer of bloodborne tracers.

### 3.5. CLDN5 level is reduced in the SVZ of fetal brains in high-dose IAV only

To determine if IAV infection during pregnancy may induce vascular barrier dysfunction in the fetal brain, we assessed gene expression and immunofluorescence of key vascular endothelial TJPs, as we did for the placenta. Given CLDN5’s critical role in brain development during gestation (Whish et al. 2015; Nitta et al. 2003) and its status as the most highly expressed TJP in the brain (Greene, Hanley, and Campbell 2019), we focused on this protein. Representative images of CD31 and CLDN5 staining in the fetal brain are shown in **Fig. 5A** (SVZ) and **Supp. Fig. S4A** (whole sagittal brain sections). Quantitative analysis of fluorescence intensity across the whole brain revealed no statistically significant differences in CD31 MFI/mm² (**Supp. Fig. S4B**) or CLDN5 (**Supp. Fig. S4C**) across treatment groups. Similarly, gene expression analysis of TJPs (**Fig. 5B** & **Supp. Fig. S4D**-**H**) and inflammatory mediators (**Supp. Fig. S4I-J**) in whole fetal brains showed no significant alterations between groups, except for an upregulation of *Cldn1* mRNA levels observed exclusively in the X31_mod_ group (**Supp. Fig. S4D**). Interestingly, mRNA transcripts for *Cldn1* and *Cldn4* were numerically reduced in both the X31_mod_ and X31_hi_ groups, but this did not reach significance (p = 0.08 and p = 0.07, respectively; **Supp. Fig. S4E-F**), likely due to variability at this embryonic stage.

**Figure 5.**
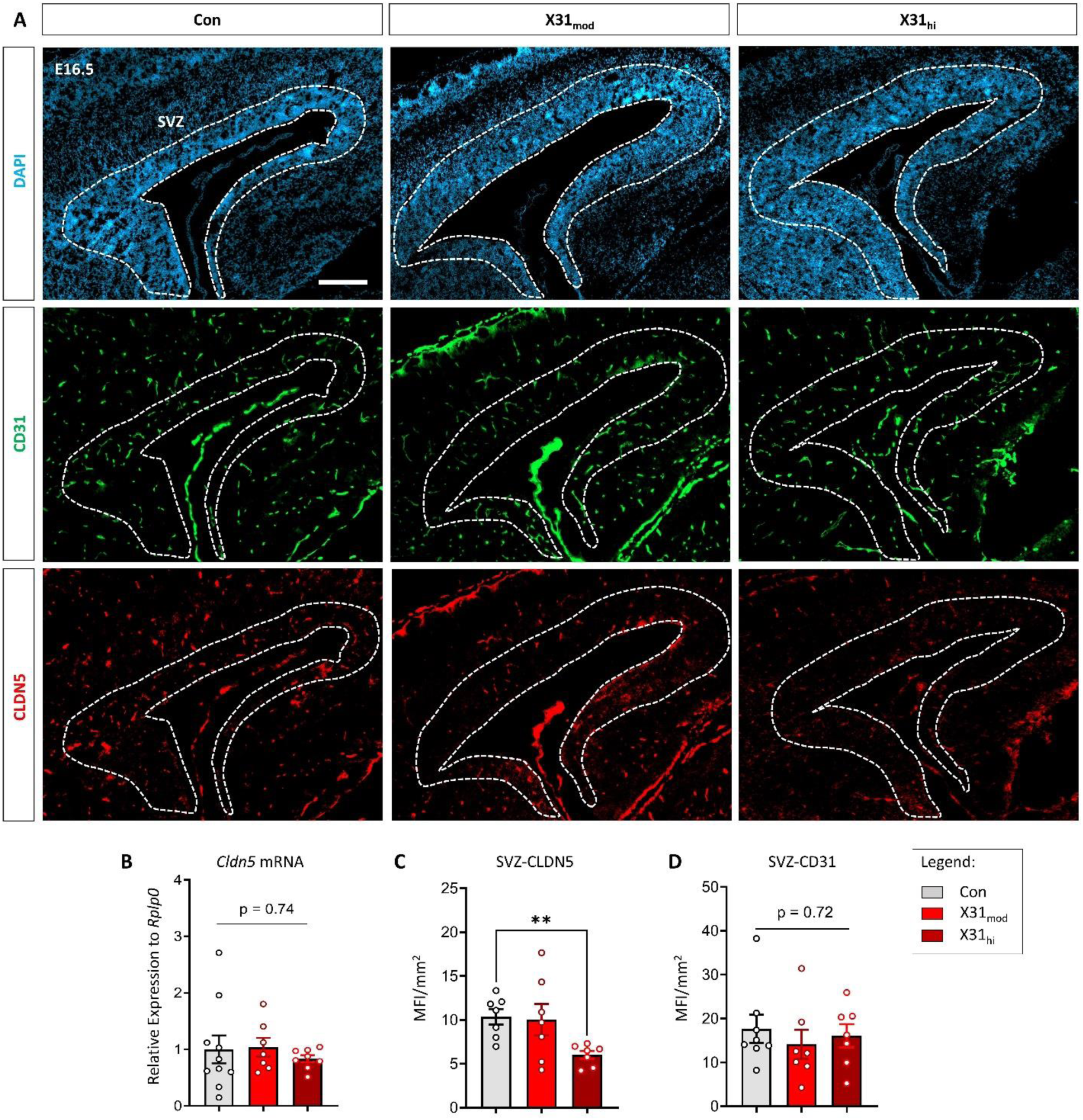
Gestational IAV infection reduces CLDN5 abundance in the fetal lateral ventricle SVZ in a dose-dependent manner. (**A**) Representative sagittal cross-section of DAPI, CLDN5, and CD31 staining in fetal brain samples at E16.5. (**B**) No significant changes were detected in CLDN5 gene expression in fetal whole brains. (**C**) MFI/mm^2^ of CLDN5 at the SVZ was reduced in fetuses exposed to X31_hi_ compared to the control group. (**D**) No changes were seen in CD31 MFI/mm^2^ across groups. Statistical analysis was conducted using a one-way ANOVA with Tukey’s post hoc test. For data containing residuals with unequal variance, Brown-Forsythe and Welch’s ANOVA with Dunnett T3 post hoc multiple comparisons was used. ** = p < 0.01. Data are expressed as means ± SEM; circles represent individual fetuses, with one representative fetus per litter; n per group: Con = 7-10, X31_mod_ = 7, X31_hi_ = 7-8. SVZ was delineated (white dots) based on DAPI staining and using reference brain atlases (Schambra 2008; GENSAT, n.d.; Allen Institute for Brain Science 2004; Chen et al. 2017). Scale bar = 200 μm. SVZ = subventricular zone.

In contrast, while TJP expression remained unchanged in whole fetal brain samples, region-specific differences were evident in the SVZ of the lateral ventricle, particularly near the ChP. A pronounced reduction in CLDN5 MFI/mm² was observed exclusively in fetuses from the X31_hi_ group (**Fig. 5C**), whereas CD31 MFI/mm² remained consistent across treatment groups (**Fig. 5D**). This decrease in CLDN5 immunofluorescence within the SVZ may represent a potential breakdown in BBB integrity, facilitating the entry of pro-inflammatory mediators from circulation into the fetal brain parenchyma.

### 3.6. Increased fibrinogen in the SVZ and ChP of the fetal brain correlates with heightened immunofluorescence of Iba1^+^ cells following maternal high-dose IAV infection

Based on the observed reduction in CLDN5 MFI/mm2 at the SVZ, we next examined the potential transfer of the bloodborne glycoprotein fibrinogen at this brain region (**Fig. 6A**). Fibrinogen has recently garnered attention due to its involvement in several neuroinflammatory conditions (Davalos and Akassoglou 2012; Mendiola et al. 2023; Petersen, Ryu, and Akassoglou 2018). In MS, AD, and TBI, fibrinogen can leak from the bloodstream into the brain parenchyma where it interacts with microglia after its cleavage to fibrin (Ryu et al. 2015; Davalos et al. 2012; Merlini et al. 2019). This interaction initiates the subsequent release of pro-inflammatory cytokines and ROS from microglia, exacerbating the disease state in adult brains (Dean et al. 2024; Roseborough et al. 2023). In these contexts, fibrinogen can also directly compromise BBB functionality, contributing to neuronal damage and exacerbating neuroinflammation (Alruwaili et al. 2023a; McLarnon 2021). Notably, fibrinogen deposition specifically at the SVZ has been demonstrated in an adult TBI model (Pous et al. 2020). To determine whether similar mechanisms may occur in the fetal brain following maternal IAV infection, we examined fibrinogen and Iba1 macrophage/microglia immunofluorescence at the SVZ and ChP within the lateral ventricle.

**Figure 6.**
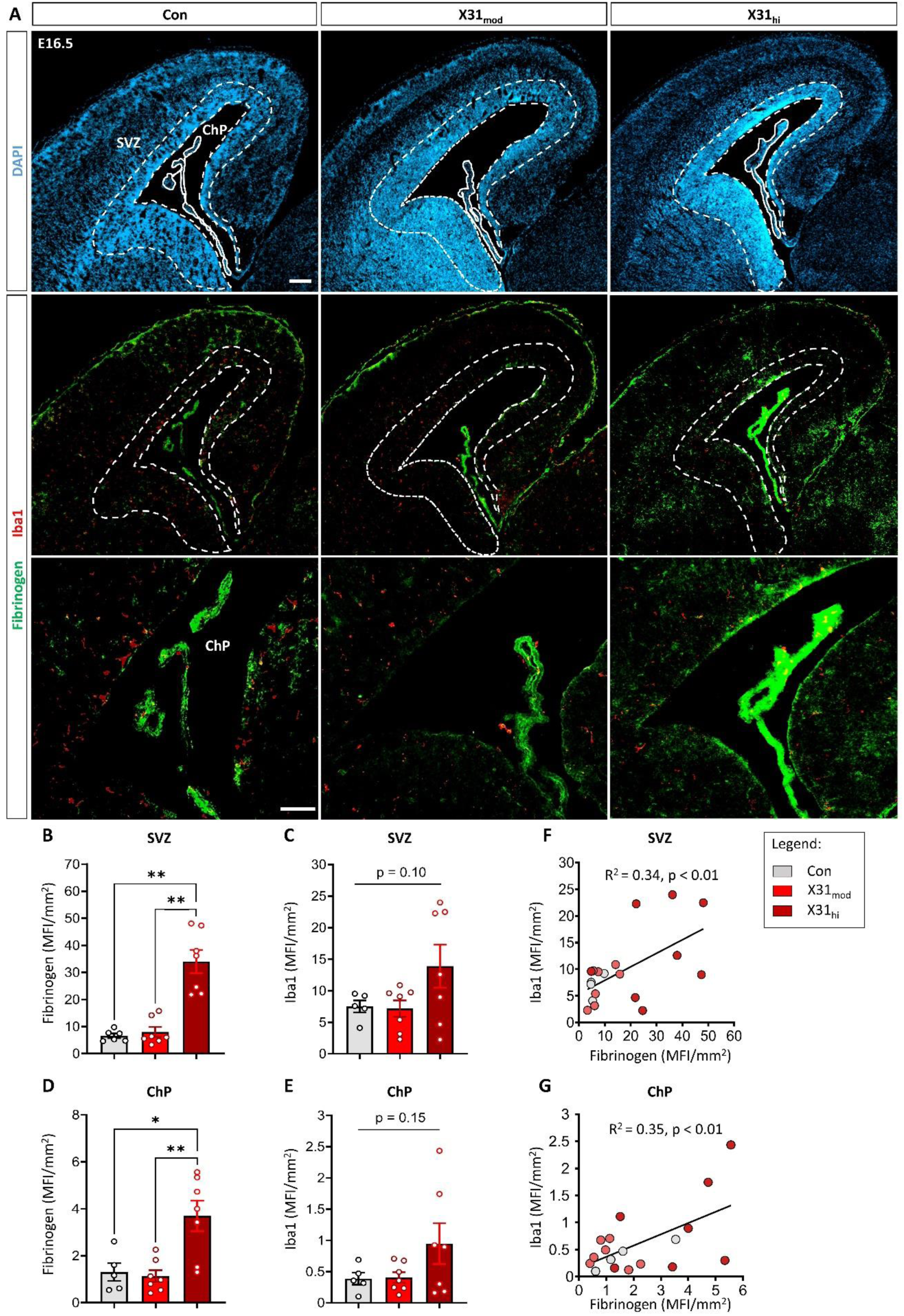
Gestational IAV infection increases fibrinogen accumulation in the SVZ and ChP, which correlates with macrophage/microglia marker Iba1. (**A**) Representative cross-sectional images depicting DAPI, fibrinogen, and Iba1 staining in fetal brain samples across groups. (**B** & **D**) Fibrinogen immunofluorescence quantified in the SVZ and ChP of the lateral ventricle was increased in the X31_hi_ group only. (**C** & **E**) MFI/mm² of Iba1, while numerically increased in the X31_hi_ group, did not reach significance within the same regions of interest. (**F-G**) Pearson correlation coefficient between fibrinogen and Iba1 MFI/mm^2^ indicates a positive correlation in both the SVZ and the ChP. Values were compared across groups using one-way ANOVA with Tukey post hoc for multiple comparisons. A Brown-Forsythe ANOVA, with Dunnett’s T3 correction for multiple comparisons, was used to analyze data with unequal variance. When residuals did not meet the assumptions of normality or homogeneity of variance, a Kruskal-Wallis ANOVA was conducted with Dunn’s post hoc correction to account for multiple comparisons; * = p < 0.05, ** = p < 0.01. Data are expressed as means ± SEM; circles represent individual fetuses, with one representative fetus per litter; n per group: Con = 4-6, X31_mod_ = 6-7, X31_hi_ = 7. Non-significant p-values were obtained from group comparisons using one-way ANOVA. SVZ outlined (white dots) using DAPI staining based on(Allen Institute for Brain Science 2004; Schambra 2008; GENSAT, n.d.; Chen et al. 2017). ChP was outlined with white solid lines (Scale bar = 200 μm). The bottom row depicts magnified ChP (bottom images, scale bar = 100 μm). SVZ = subventricular zone; ChP = choroid plexus.

Indeed, we observed an increase in fibrinogen MFI levels in both the SVZ (**Fig. 6B**) and ChP (**Fig. 6D**) only in fetal brains from the X31_hi_ group. While a numerical increase in SVZ and ChP Iba1 MFI/mm^2^ in the X31_hi_ group did not reach significance (p = 0.10 and p = 0.15, respectively; **Fig. 6C** & **E**), overall number of Iba1^+^ cells remained relatively constant (**Supp. Fig. S5A** & **C**). Still, a positive correlation was observed between fibrinogen MFI/mm² and Iba1 MFI/mm² in both brain regions (**Fig. 6F** & **G**). Iba1^+^ cell counts in the SVZ and ChP (**Supp. Fig. S5B** & **D**) did not correlate with fibrinogen MFI/mm². Taken together, these findings indicate that the SVZ and ChP may serve as hot spots for leakage of bloodborne molecules, including fibrinogen, during IAV-induced MIA. To our knowledge, this is the first time such a phenomenon has been observed.

## 4. Discussion

While maternal viral infection is a known risk factor for offspring NDDs, the exact mechanisms mediating this risk are still incompletely understood. Here, we characterize an underexplored feature of maternal inflammation—a breakdown in placental and fetal brain barriers—in a biologically relevant mouse model of live respiratory IAV infection. Consistent with our previous reports (Antonson et al. 2021), the severity of barrier breakdown was dose-dependent, further bolstering the existence of an infection severity threshold that mimics epidemiological data. Unsurprisingly, infection-induced increases in transplacental transfer of bloodborne molecules appeared to preferentially impact brain areas known to have more permeable barriers, like the SVZ and ChP. To our knowledge, this is the first time fibrinogen accumulation in the fetal brain has been documented. Collectively, our findings reinforce the hypothesis that the SVZ and ChP serve as entry points for the transfer of bloodborne molecules during maternal inflammatory insult. Although this hypothesis warrants further investigation, we believe that our findings offer compelling initial evidence for IAV-induced leakage of bloodborne molecules that could directly mediate neurodevelopmental outcomes.

Gestational IAV infection is known to impair placental function and lead to fetal growth restriction (Cotechini, Hopman, and Graham 2014; D. X. Xu et al. 2006; Creisher et al. 2024; Liong et al. 2020; Otero et al. 2024; Antonson et al. 2021)—adverse outcomes that have been associated with increased risk for NDDs in offspring (Van Campen et al. 2020; Jiao Wang et al. 2024; Oseghale et al. 2022). Using the same mouse-adapted IAV strain at a dose comparable to our X31_hi_ group, Liong et al. (2020) demonstrated that maternal IAV infection impairs fetal and placental growth. They documented a pronounced elevation in proinflammatory mediators, leading to a “vascular storm” that caused systemic endothelial dysfunction in major maternal arteries. Consequently, placental blood flow—and thus oxygen supply to the fetal brain—was significantly impaired (Liong et al. 2020). Our observations of dose-dependent intrauterine growth restriction complement these findings, indicating that a threshold of infection severity must be exceeded to elicit significant restrictions in placental mass and fetal size. However, even severe IAV insult was not sufficient to induce fetal loss. Notably, viral transcript levels in the lung did not differ between the two doses, suggesting that host immune responses, rather than direct viral burden, are the primary drivers of growth impairments. However, because our analysis was limited to a single gestational time point (7 dpi, GD16.5), the full trajectory of immune-mediated effects across pregnancy requires further exploration.

We hypothesized that systemic anti-viral inflammatory responses may compromise the integrity and functionality of the maternal-fetal barrier, enabling the uncontrolled passage of pro-inflammatory mediators into the fetal compartment. Our previous studies provided evidence of placental pathology, including necrosis, mineralization, and neutrophil infiltration, following maternal exposure to a moderate dose of X31 (Antonson et al. 2021). In line with this, other rodent studies have shown that mimetic-induced MIA (e.g., lipopolysaccharide or poly I:C) disrupts placental structure and impairs fetal BBB development, leading to persistent dysfunction and chronic neuroinflammation (Simões, Sangiogo, et al. 2018; Zhao et al. 2022). Collectively, these studies support the notion that placental impairment—particularly disruptions to vascular integrity and blood flow—may enable the transfer of maternally derived agents, typically excluded from the *in utero* environment, into the fetal compartment. This influx of bloodborne molecules could adversely affect neurodevelopment in part through direct compromise of fetal BBB formation and function. While we did not observe significant alterations in TJP expression or CD31 immunofluorescence, these measurements may not fully capture the extent of placental dysfunction. Importantly, placental malperfusion— such as the vascular pathology we previously reported (Antonson et al. 2021) in X31_mod_ pregnancies—may occur independently of molecular or immunohistochemical changes typically associated with barrier integrity. As such, even in the absence of overt changes in gene expression or protein localization, underlying vascular abnormalities may represent a parallel mechanism by which maternal inflammation impairs placental function, thereby permitting aberrant passage of pro-inflammatory mediators into the fetal milieu. To address this possibility, targeted assessments of placental vascular pathology that extend beyond traditional markers of barrier dynamics and integrity are warranted.

Here, we observed an increase in the transplacental transfer of three maternally derived relatively high molecular weight tracers following IAV infection. Interestingly, the distinct patterns of tracer distribution across placental layers suggest that MIA may differentially affect placental permeability depending on the severity of the immune challenge. In the X31_mod_ group, the reduction in Cy5 and Fluorescein MFI—localized primarily to the labyrinth—could indicate a functional alteration in molecular transport or retention in this region, where there is maximal maternal-fetal blood exchange. Meanwhile, the significant accumulation of TxRed and Fluorescein in the junctional zone of the X31_hi_ group points toward increased tracer passage or reduced clearance. This potentially reflects a breakdown in region-specific regulation under heightened inflammatory conditions. The absence of changes in the decidua highlights the specificity of these effects. Together, these findings support the notion that MIA does not uniformly disrupt placental integrity but rather induces region-specific changes that may facilitate abnormal transplacental transfer and influence fetal exposure to maternal signals. Extending beyond the placenta, tracer accumulation was most prominent in the fetal liver, reinforcing its role as a central site for processing maternally derived bloodborne molecules, particularly under inflammatory conditions (R. Xu et al. 2014; Jing Wang et al. 2017; Jaeschke, Farhood, and Smith 1990).

Beyond systemic distribution, our region-specific analysis of the fetal brain revealed a discrete accumulation of 500 KDa Fluorescein tracer within the SVZ of the lateral ventricle, which coincided with a decrease in CLDN5 immunofluorescence in this same region. Given the critical role of CLDN5 in maintaining BBB integrity (Hashimoto et al. 2023; Berndt et al. 2019; Nitta et al. 2003), its reduction in the SVZ may indicate a local compromise in barrier function, rendering this region particularly susceptible to the influx of peripheral molecules. The SVZ naturally exhibits greater permeability due to its lower pericyte coverage and specialized vasculature (Tavazoie et al. 2008), which could further contribute to its heightened vulnerability during inflammatory conditions. Interestingly, there appeared to be preferential accumulation of Fluorescein but not the smaller TxRed and Cy5 tracers (70 and 250 KDa, respectively). This pattern may reflect differential kinetics of tracer retention and clearance within the brain parenchyma. The smaller tracers, due to their lower molecular weight, are more likely to penetrate deeper into the brain tissue and undergo rapid diffusion and washout. Thus, local detectability of relatively smaller tracers may be reduced in specific regions like the SVZ. In contrast, the larger 500 kDa Fluorescein tracer may exhibit slower parenchymal penetration and reduced clearance, resulting in greater localized retention and enhanced signal at the site of barrier disruption. Although our analysis focused on the SVZ, assessing additional neuroanatomical sites would be valuable to determine if similar BBB disruptions occur elsewhere in the developing brain. Nevertheless, our findings suggest that maternal IAV exposure leads to dose- and region-specific alterations in BBB permeability, facilitating the entry of molecular mediators from both fetal and maternal circulation into the brain parenchyma.

Moreover, previous studies indicate that inflammatory mediators in circulation (e.g., TNF-α, IL-1β, IL-6, and IFN-γ) can modulate TJP expression, potentially exacerbating BBB permeability (Huang, Hussain, and Chang 2021; Varatharaj and Galea 2017; Galea 2021; TSAO et al. 2001; Blamire et al. 2000). In the context of MIA, these inflammatory molecules may compromise vascular endothelial barrier integrity, leading to increased BBB permeability (Pearson and Iadecola 2022; Mohebalizadeh et al. 2024; Simões, Sangiogo, et al. 2018; Simões, Generoso, et al. 2018). Such localized permeability shifts could be particularly pronounced in regions like the SVZ, aligning with our findings of reduced CLDN5 MFI. These observations raise critical questions about whether these changes in permeability are transient, which might reflect an acute inflammatory response, or if they have lasting consequences for fetal neurodevelopment by chronically altering the local microenvironment (Zhao et al. 2022).

Considering these questions, it is essential to identify which circulating factors might exacerbate changes in the brain’s proinflammatory state. One prominent candidate is the bloodborne protein fibrinogen. While this molecule is essential for vascular healing following injury (Pieters and Wolberg 2019), it has also been shown to enhance vascular permeability at the BBB and amplify the oxidative immune response under neuroinflammatory conditions (Muradashvili et al. 2011; Guo et al. 2009; Patibandla et al. 2009; Wood 2019). In models of neuroinflammatory disorders in both humans and rodents, disruption of the BBB has been shown to permit the extravasation of fibrinogen from the vascular lumen into the brain parenchyma. Within neural tissue, fibrinogen is converted into a range of bioactive byproducts, most notably fibrin, which functions as a key effector molecule for the initiation of inflammatory and pro-oxidative cascade (Alruwaili et al. 2023b; Ahn et al. 2019). Notably, a recent study in a mouse model of TBI-induced neuroinflammation demonstrated significant fibrinogen spread that was particularly evident in the SVZ (Pous et al. 2020). Our observation of a robust dose-dependent accumulation of fibrinogen in the fetal SVZ and ChP following maternal IAV infection indicates that similar region-specific patterns of inflammation-induced vascular breakdown may also occur prenatally. Furthermore, this finding supports our hypothesis that the SVZ may serve as a potential entry point for a multitude of circulating proinflammatory agents in our IAV model.

Importantly, fibrinogen accumulation correlated with increased Iba1^+^ microglia/macrophage immunofluorescence, suggesting that leaked fibrinogen may be sensed by these cells, triggering pro-oxidative responses. In adults, microglia detect cleaved fibrinogen (fibrin) via the CD11b/CD18 (Mac-1) integrin receptor, activating NADPH oxidase and generating ROS, which amplifies oxidative stress and neuroinflammation (Zhang et al. 2021; Kračun et al. 2025; Simpson and Oliver 2020; Wen and Zhang 2023). Fibrinogen also directly impairs neurite outgrowth through β3 integrin and EGFR-dependent signaling (Schachtrup et al. 2007b), disrupting cytoskeletal remodeling essential for axon extension. In the developing brain, this dual mechanism may compromise normal brain development (Weaver et al. 2024). The observed spatial overlap between fibrinogen and Iba1^+^ cells in our study supports the possibility that fetal microglia respond similarly to fibrin by producing ROS, which, alongside fibrin’s direct effects on neurons, may contribute to neurodevelopmental injury. This is particularly concerning given the pro-neurogenic role of fetal microglia during brain development (Luo and Wang 2024; Michell-Robinson et al. 2015; Tong and Vidyadaran 2016; Mastenbroek et al. 2024). MIA-induced shifts in microglial responses have been shown to disrupt key signaling pathways essential for neuronal growth, axonal and dendritic branching, and synapse formation (Matelski et al. 2021; Lee et al. 2022; Györffy et al. 2016; Li and Barres 2018; Thion et al. 2019). Fibrinogen/fibrin deposition in the brain parenchyma during IAV infection may prime fetal microglia/macrophages, promoting neuroinflammatory and oxidative stress responses that contribute to neuropathology. Future and ongoing investigations by our group are focused on answering this question.

While emerging evidence increasingly highlights sex-specific effects of MIA on microglial function and BBB integrity (Hui et al. 2020; Rasile et al. 2022; Weber and Clyne 2021), our current study was not designed to evaluate such differences. Therefore, future investigations exploring sex-dependent effects in this model will be essential for a more complete understanding of how maternal infection may differentially impact brain development in males and females.

Overall, our study provides direct evidence that maternal IAV infection enhances transplacental transfer of bloodborne molecules into key regions of the fetal brain—the SVZ and ChP in a dose-dependent manner. The increase in fibrinogen levels, and the correlation with Iba1 in these areas, underscores a potential link between infection-induced barrier permeability, fibrinogen deposition, and neuroinflammatory processes in the developing brain. These findings highlight the importance of the fetal brain microenvironment in mediating the effects of maternal immune challenges and offer new directions for exploring novel neuroimmune mechanisms underlying NDD etiologies.

## 6. Acknowledgements

We would like to thank Dr. Katerina Akassoglou & Dr. Jae Kyu Ryu (Gladstone Institutes-University of California San Fransisco) for providing us with Sheep Anti-Fibrinogen antibody, along with invaluable technical expertise and guidance on fibrinogen-related studies. This study was supported by the Roy J. Carver Charitable Trust (Grant #23-5683), Department of Animal Sciences Matchstick Grant, and by start-up funds from the University of Illinois Urbana-Champaign to AMA.

## 7. Author Contributions

**Rafael J. Gonzalez-Ricon**: Conceptualization, Methodology, Validation, Formal Analysis, Investigation, Writing – Original Draft, Writing – Review & Editing, Visualization, Project Administration. **Ashley M. Otero**: Investigation, Writing – Review & Editing. **Izan Chalen**: Investigation, Review & Editing. **Jeffery N. Savas**: Review & Editing. **Adrienne M. Antonson**: Conceptualization, Methodology, Writing – Review & Editing, Supervision, Funding Acquisition.

