## Supplementary Figures & Tables for "Influenza A virus infection during pregnancy increases transfer of maternal bloodborne molecules to fetal tissues"

### Slide 1
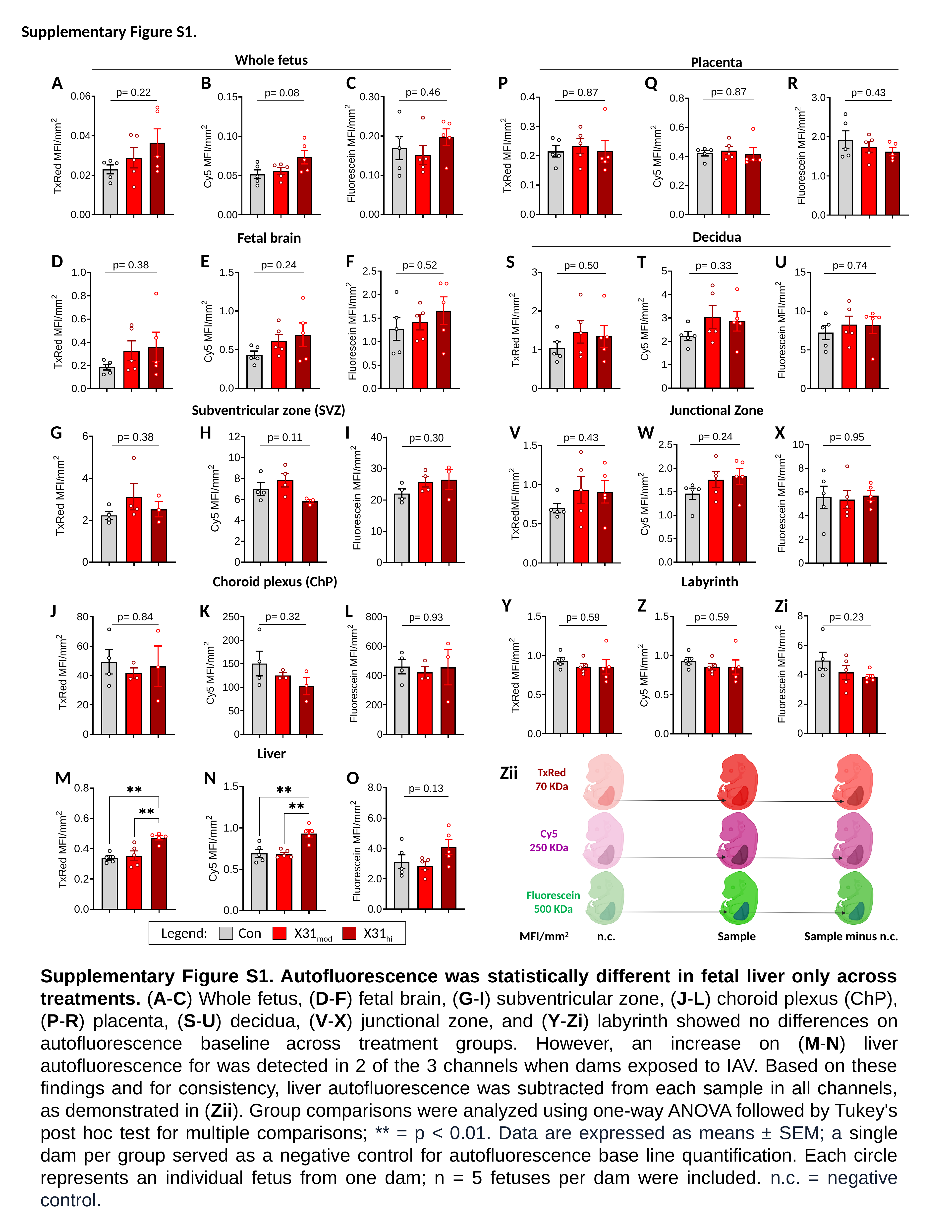

Supplementary Figure S1.
Whole fetus
Placenta
A
B
C
P
Q
R
Decidua
Fetal brain
D
E
F
S
T
U
Subventricular zone (SVZ)
Junctional Zone
G
H
I
V
W
X
Choroid plexus (ChP)
Labyrinth
Y
Z
Zi
J
K
L
Liver
TxRed
70 KDa
Cy5
250 KDa
Fluorescein
500 KDa
 Sample
 Sample minus n.c.
MFI/mm^2^
n.c.
Zii
M
N
O
Legend:
X31_hi_
X31_mod_
Con
Supplementary Figure S1. Autofluorescence was statistically different in fetal liver only across treatments. (A-C) Whole fetus, (D-F) fetal brain, (G-I) subventricular zone, (J-L) choroid plexus (ChP), (P-R) placenta, (S-U) decidua, (V-X) junctional zone, and (Y-Zi) labyrinth showed no differences on autofluorescence baseline across treatment groups. However, an increase on (M-N) liver autofluorescence for was detected in 2 of the 3 channels when dams exposed to IAV. Based on these findings and for consistency, liver autofluorescence was subtracted from each sample in all channels, as demonstrated in (Zii). Group comparisons were analyzed using one-way ANOVA followed by Tukey's post hoc test for multiple comparisons; ** = p < 0.01. Data are expressed as means ± SEM; a single dam per group served as a negative control for autofluorescence base line quantification. Each circle represents an individual fetus from one dam; n = 5 fetuses per dam were included. n.c. = negative control.

### Slide 2
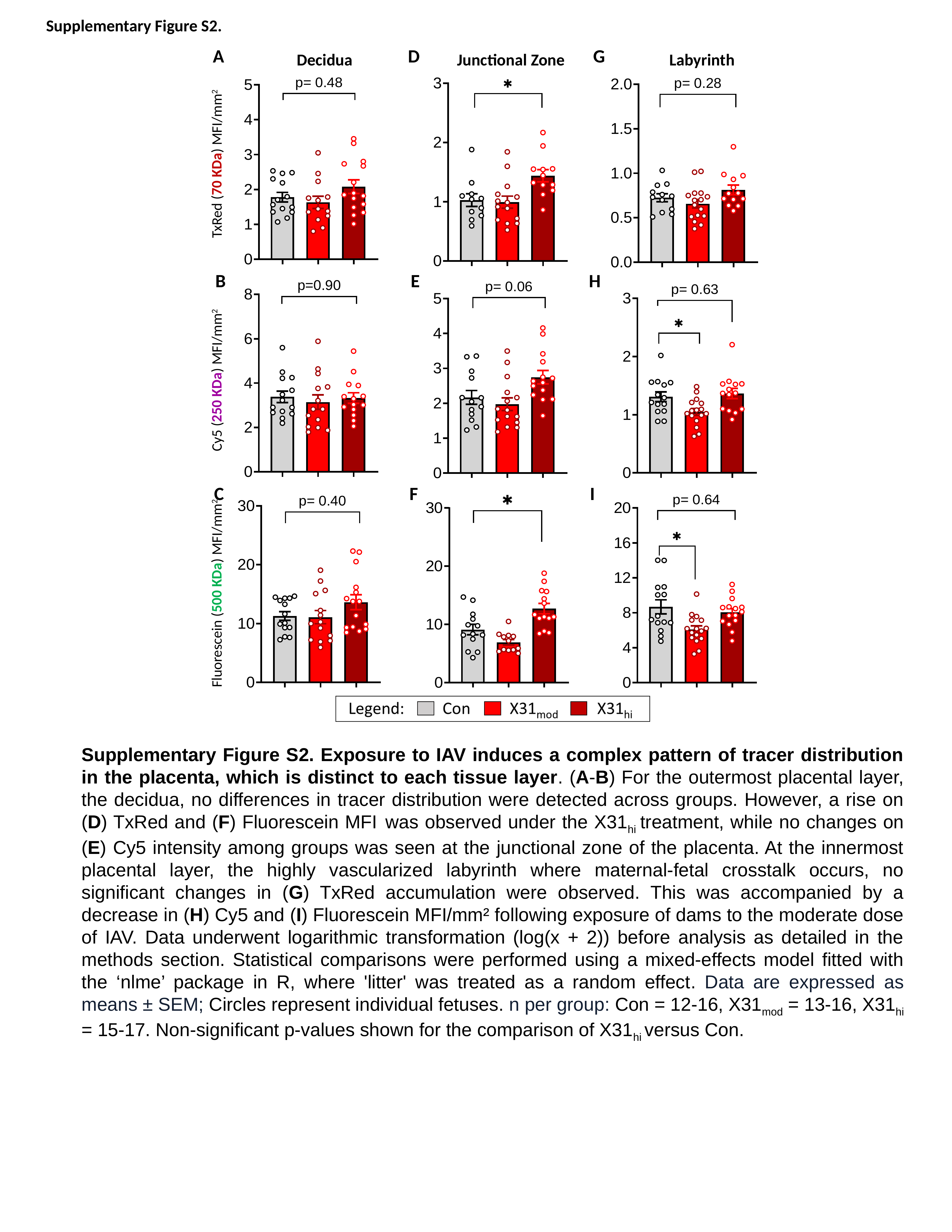

Supplementary Figure S2.
A
D
G
 Decidua Junctional Zone Labyrinth
TxRed (70 KDa) MFI/mm^2^
B
E
H
Cy5 (250 KDa) MFI/mm^2^
C
F
I
Fluorescein (500 KDa) MFI/mm^2^
Supplementary Figure S2. Exposure to IAV induces a complex pattern of tracer distribution in the placenta, which is distinct to each tissue layer. (A-B) For the outermost placental layer, the decidua, no differences in tracer distribution were detected across groups. However, a rise on (D) TxRed and (F) Fluorescein MFI was observed under the X31_hi_ treatment, while no changes on (E) Cy5 intensity among groups was seen at the junctional zone of the placenta. At the innermost placental layer, the highly vascularized labyrinth where maternal-fetal crosstalk occurs, no significant changes in (G) TxRed accumulation were observed. This was accompanied by a decrease in (H) Cy5 and (I) Fluorescein MFI/mm² following exposure of dams to the moderate dose of IAV. Data underwent logarithmic transformation (log(x + 2)) before analysis as detailed in the methods section. Statistical comparisons were performed using a mixed-effects model fitted with the ‘nlme’ package in R, where 'litter' was treated as a random effect. Data are expressed as means ± SEM; Circles represent individual fetuses. n per group: Con = 12-16, X31_mod_ = 13-16, X31_hi_ = 15-17. Non-significant p-values shown for the comparison of X31_hi_ versus Con.

### Slide 3
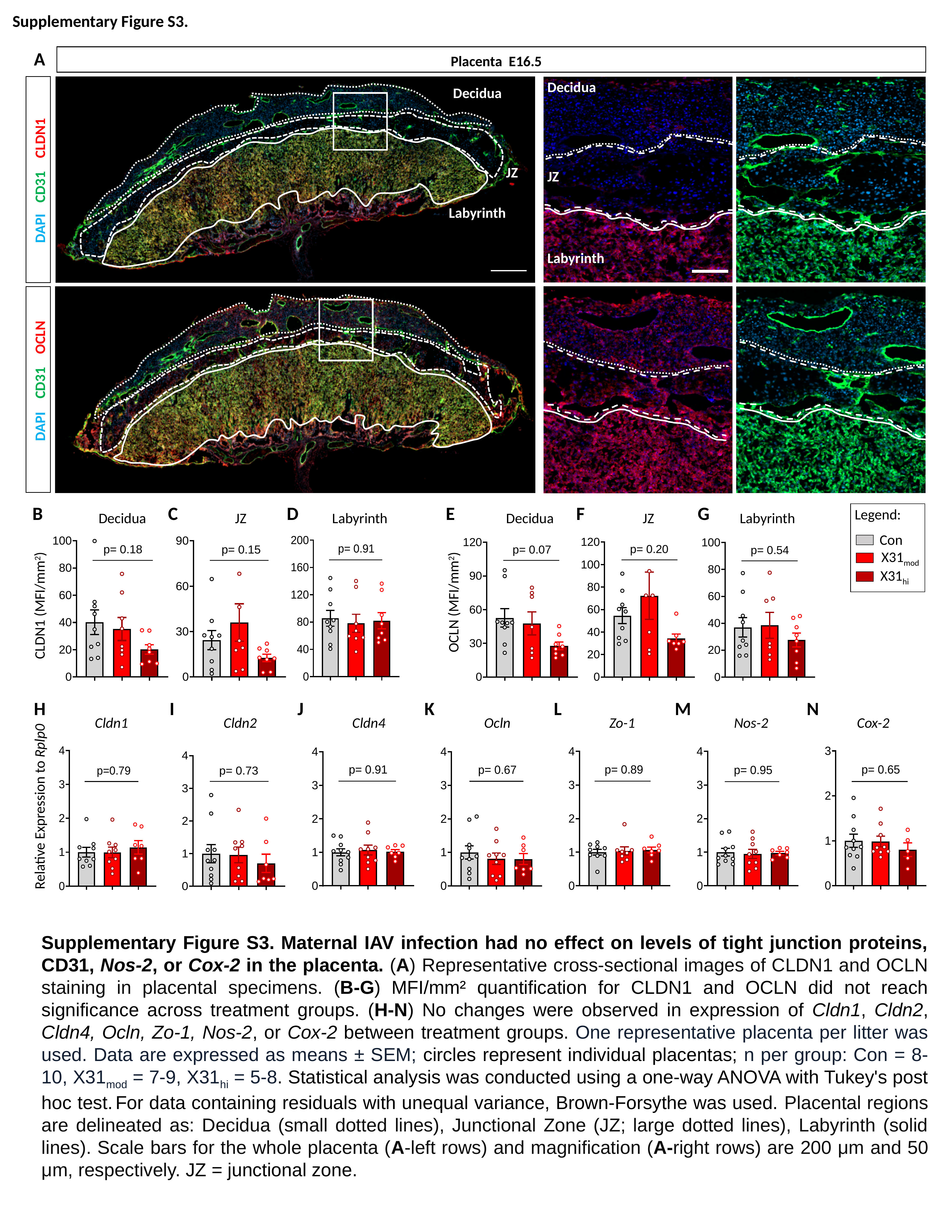

Supplementary Figure S3.
A
Placenta E16.5
Decidua
Decidua
CLDN1
CD31
DAPI
OCLN
CD31
DAPI
JZ
JZ
Labyrinth
Labyrinth
B
C
G
D
E
F
Legend:
Con
X31_mod_
X31_hi_
 Decidua JZ Labyrinth Decidua JZ Labyrinth
OCLN (MFI/mm^2^)
CLDN1 (MFI/mm^2^)
H
I
M
N
J
K
L
 Cldn1 Cldn2 Cldn4 Ocln Zo-1 Nos-2 Cox-2
Relative Expression to Rplp0
Supplementary Figure S3. Maternal IAV infection had no effect on levels of tight junction proteins, CD31, Nos-2, or Cox-2 in the placenta. (A) Representative cross-sectional images of CLDN1 and OCLN staining in placental specimens. (B-G) MFI/mm² quantification for CLDN1 and OCLN did not reach significance across treatment groups. (H-N) No changes were observed in expression of Cldn1, Cldn2, Cldn4, Ocln, Zo-1, Nos-2, or Cox-2 between treatment groups. One representative placenta per litter was used. Data are expressed as means ± SEM; circles represent individual placentas; n per group: Con = 8-10, X31_mod_ = 7-9, X31_hi_ = 5-8. Statistical analysis was conducted using a one-way ANOVA with Tukey's post hoc test. For data containing residuals with unequal variance, Brown-Forsythe was used. Placental regions are delineated as: Decidua (small dotted lines), Junctional Zone (JZ; large dotted lines), Labyrinth (solid lines). Scale bars for the whole placenta (A-left rows) and magnification (A-right rows) are 200 μm and 50 μm, respectively. JZ = junctional zone.

### Slide 4
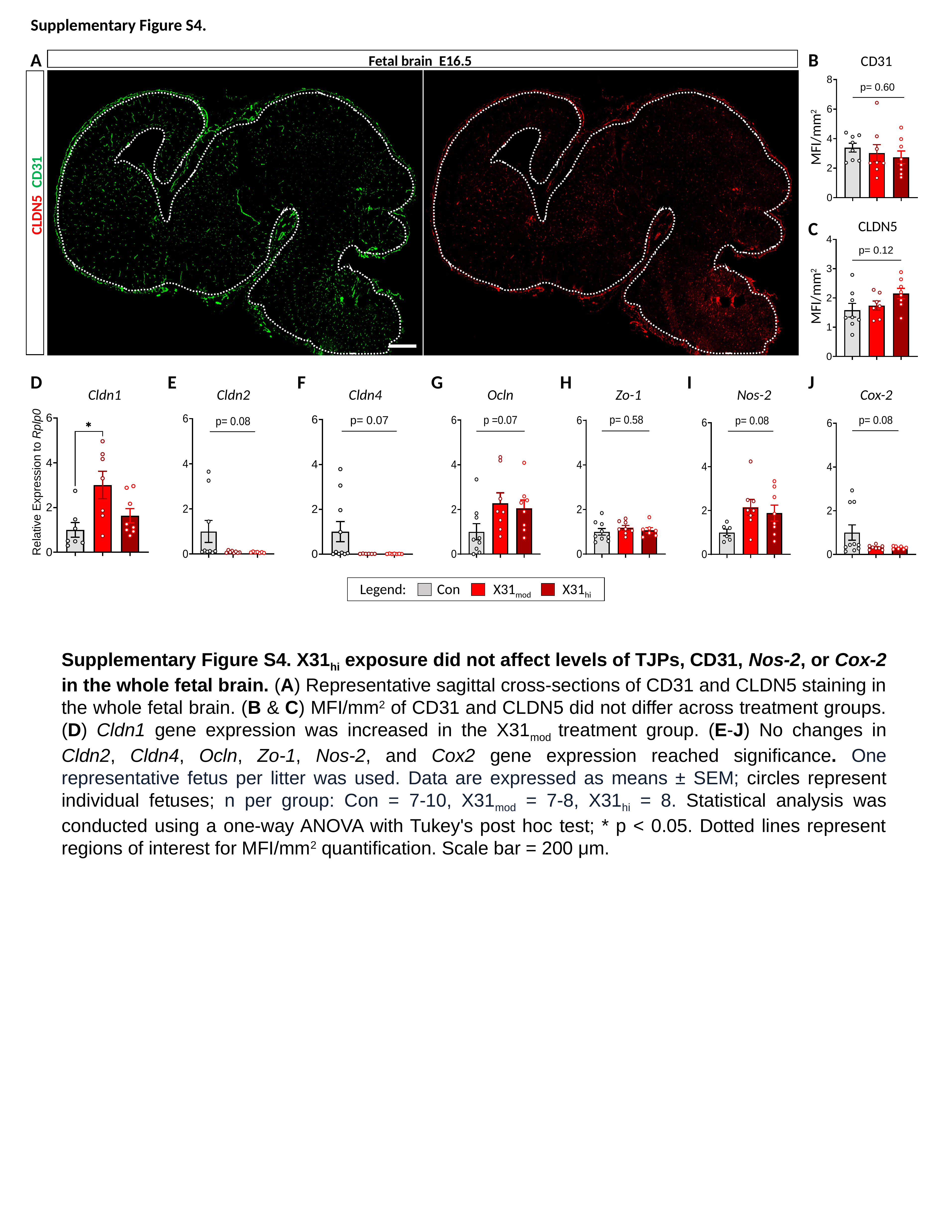

Supplementary Figure S4.
A
B
Fetal brain E16.5
CD31
MFI/mm^2^
CD31
CLDN5
C
CLDN5
MFI/mm^2^
D
E
I
J
F
G
H
 Cldn1 Cldn2 Cldn4 Ocln Zo-1 Nos-2 Cox-2
Legend:
X31_hi_
X31_mod_
Con
Supplementary Figure S4. X31_hi_ exposure did not affect levels of TJPs, CD31, Nos-2, or Cox-2 in the whole fetal brain. (A) Representative sagittal cross-sections of CD31 and CLDN5 staining in the whole fetal brain. (B & C) MFI/mm^2^ of CD31 and CLDN5 did not differ across treatment groups. (D) Cldn1 gene expression was increased in the X31_mod_ treatment group. (E-J) No changes in Cldn2, Cldn4, Ocln, Zo-1, Nos-2, and Cox2 gene expression reached significance. One representative fetus per litter was used. Data are expressed as means ± SEM; circles represent individual fetuses; n per group: Con = 7-10, X31_mod_ = 7-8, X31_hi_ = 8. Statistical analysis was conducted using a one-way ANOVA with Tukey's post hoc test; * p < 0.05. Dotted lines represent regions of interest for MFI/mm^2^ quantification. Scale bar = 200 μm.

### Slide 5
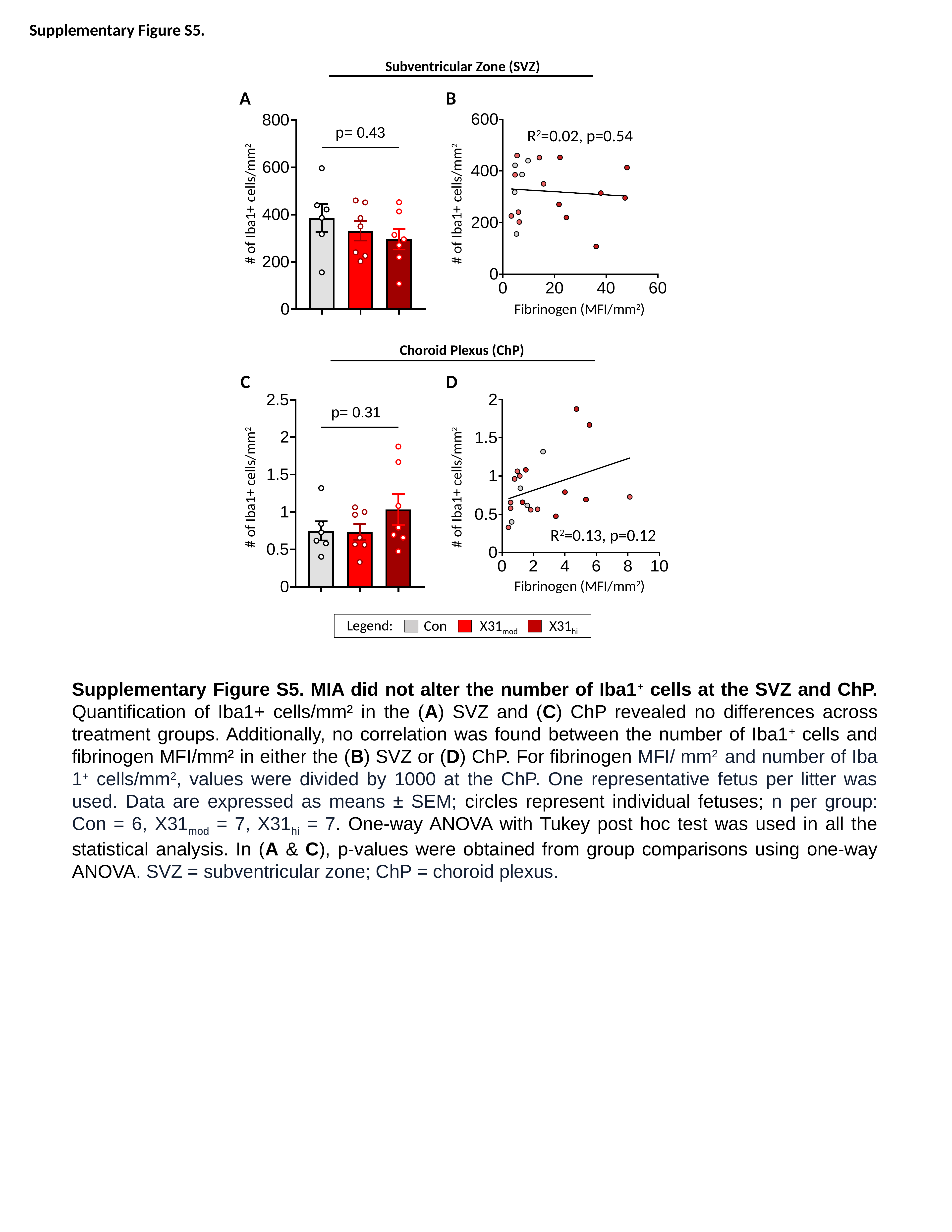

Supplementary Figure S5.
Subventricular Zone (SVZ)
A
B
R^2^=0.02, p=0.54
#### of Iba1+ cells/mm^2^
#### of Iba1+ cells/mm^2^
Fibrinogen (MFI/mm^2^)
Choroid Plexus (ChP)
C
D
#### of Iba1+ cells/mm^2^
#### of Iba1+ cells/mm^2^
R^2^=0.13, p=0.12
Fibrinogen (MFI/mm^2^)
Legend:
X31_hi_
X31_mod_
Con
Supplementary Figure S5. MIA did not alter the number of Iba1^+^ cells at the SVZ and ChP. Quantification of Iba1+ cells/mm² in the (A) SVZ and (C) ChP revealed no differences across treatment groups. Additionally, no correlation was found between the number of Iba1^+^ cells and fibrinogen MFI/mm² in either the (B) SVZ or (D) ChP. For fibrinogen MFI/ mm^2^ and number of Iba 1^+^ cells/mm^2^, values were divided by 1000 at the ChP. One representative fetus per litter was used. Data are expressed as means ± SEM; circles represent individual fetuses; n per group: Con = 6, X31_mod_ = 7, X31_hi_ = 7. One-way ANOVA with Tukey post hoc test was used in all the statistical analysis. In (A & C), p-values were obtained from group comparisons using one-way ANOVA. SVZ = subventricular zone; ChP = choroid plexus.

### Slide 6
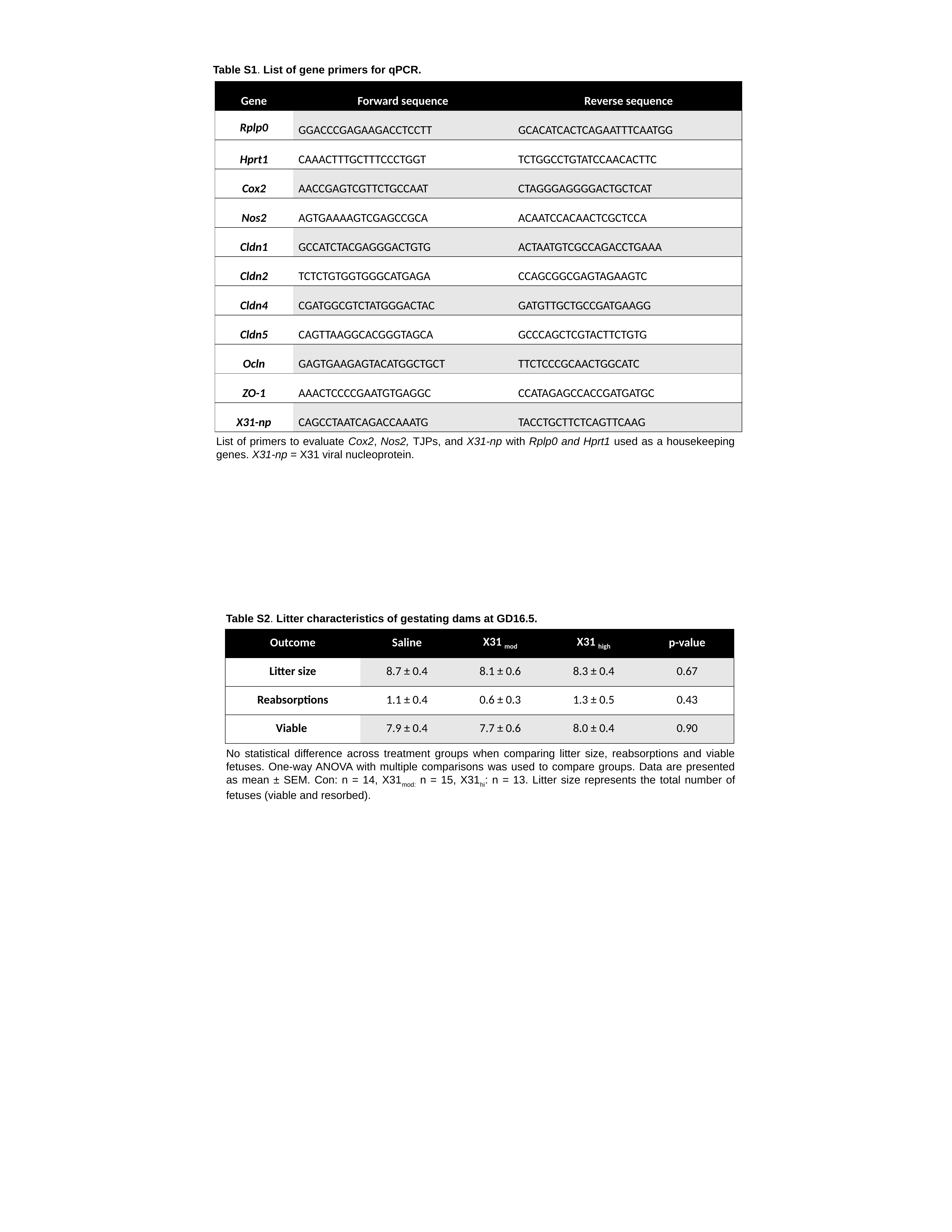

Table S1. List of gene primers for qPCR.
| Gene | Forward sequence | Reverse sequence |
| --- | --- | --- |
| Rplp0 | GGACCCGAGAAGACCTCCTT | GCACATCACTCAGAATTTCAATGG |
| Hprt1 | CAAACTTTGCTTTCCCTGGT | TCTGGCCTGTATCCAACACTTC |
| Cox2 | AACCGAGTCGTTCTGCCAAT | CTAGGGAGGGGACTGCTCAT |
| Nos2 | AGTGAAAAGTCGAGCCGCA | ACAATCCACAACTCGCTCCA |
| Cldn1 | GCCATCTACGAGGGACTGTG | ACTAATGTCGCCAGACCTGAAA |
| Cldn2 | TCTCTGTGGTGGGCATGAGA | CCAGCGGCGAGTAGAAGTC |
| Cldn4 | CGATGGCGTCTATGGGACTAC | GATGTTGCTGCCGATGAAGG |
| Cldn5 | CAGTTAAGGCACGGGTAGCA | GCCCAGCTCGTACTTCTGTG |
| Ocln | GAGTGAAGAGTACATGGCTGCT | TTCTCCCGCAACTGGCATC |
| ZO-1 | AAACTCCCCGAATGTGAGGC | CCATAGAGCCACCGATGATGC |
| X31-np | CAGCCTAATCAGACCAAATG | TACCTGCTTCTCAGTTCAAG |
List of primers to evaluate Cox2, Nos2, TJPs, and X31-np with Rplp0 and Hprt1 used as a housekeeping genes. X31-np = X31 viral nucleoprotein.
Table S2. Litter characteristics of gestating dams at GD16.5.
| Outcome | Saline | X31 _mod_ | X31 _high_ | p-value |
| --- | --- | --- | --- | --- |
| Litter size | 8.7 ± 0.4 | 8.1 ± 0.6 | 8.3 ± 0.4 | 0.67 |
| Reabsorptions | 1.1 ± 0.4 | 0.6 ± 0.3 | 1.3 ± 0.5 | 0.43 |
| Viable | 7.9 ± 0.4 | 7.7 ± 0.6 | 8.0 ± 0.4 | 0.90 |
No statistical difference across treatment groups when comparing litter size, reabsorptions and viable fetuses. One-way ANOVA with multiple comparisons was used to compare groups. Data are presented as mean ± SEM. Con: n = 14, X31_mod:_ n = 15, X31_hi_: n = 13. Litter size represents the total number of fetuses (viable and resorbed).
